# Variation in Resource Environment Drives Adaptive Divergence in *Drosophila melanogaster*

**DOI:** 10.1101/2024.12.10.626603

**Authors:** Jack K. Beltz, Mark Christopher Bitter, August Goldfischer, Dmitri A. Petrov, Paul Schmidt

**Affiliations:** University of Pennsylvania, Department of Biology 433 S University Avenue, Philadelphia, PA, 19104; Stanford University, Department of Biology 371 Jane Stanford Way, Stanford California, 94305-5020

**Keywords:** Adaptation, Seasonality, Environmental Heterogeneity, Phenotypic Evolution, Nutrition, Resource Variation, D. melanogaster, Genomic Evolution, Demographic Response

## Abstract

Natural populations often experience heterogeneity in the quality and abundance of environmentally acquired resources across both space and time, and this variation can influence population demographics and evolutionary dynamics. In this study, we directly manipulated diet in replicate populations of *Drosophila melanogaster* cultured in experimental mesocosms in the field. We found no significant effect of resource variation on demographic patterns. Furthermore, while resource variation altered the patterns of phenotypic and genomic evolution, this effect is secondary to population responses to seasonally fluctuating selective pressures. Seasonal adaptation was observed for all traits assayed and elicited genome-wide signatures of selection; in contrast, adaptation to the resource environment was trait-specific and exhibited an oligogenic architecture. This illustrates the capacity of populations to adapt to a specific axis of variation (the resource environment) without hindering the adaptive response to seasonal change. This in turn, suggests that resource variation may be an important force driving fluctuating selection across natural populations, ultimately contributing to the maintenance of genetic and phenotypic variation.

## INTRODUCTION

Identifying the environmental factors that shape population demographics and evolutionary dynamics remains a fundamental goal in ecology and evolution. A major component of variation in natural habitats is the availability and quality of resources (e.g., water, light, nitrogen, food supply, etc.). Variations in the quality or availability of critical resources can impact the growth and productivity of populations (Schindler, 1974; Blaustein *et al*., 2003; Gilg *et al*., 2003) and alter selection pressures that shift patterns of adaptive evolution (Grant & Grant, 1989). As ecological and evolutionary processes can occur on the same timescales (Johnston & Selander, 1964; Reznick *et al*., 1997; Hendry & Kinnison, 1999; Hendry *et al*., 2007), an open question is whether variation in environmental resources can simultaneously drive both ecological and evolutionary dynamics.

Environmental quality affects population demographics in various systems and contexts (MacArthur, 1984; Philippi *et al*., 1987; Pimm *et al*., 1988; Klok & De Roos, 1998; Belovsky *et al*., 1999; Hakoyama *et al*., 2000; J. S. Bancroft, 2003; Franken & Hik, 2004; Griffen & Drake, 2008). Furthermore, the quality and availability of environmentally acquired resources are used as predictors when estimating the growth rate and carrying capacity of populations (Hobbs *et al*., 1982) and may also affect different components of fitness (Mitrovski & Hoffmann, 2001; Chatelain *et al*., 2021). Resource-limited population growth is especially prevalent in systems where water or food supplies are seasonal or ephemeral, impacting local rates of extinction (Schindler, 1974; Longman *et al*., 2023).

Adaptive responses to variation in the biotic and abiotic environment can occur over extremely short intervals (Grainger *et al*., 2021; Rudman *et al*., 2022; Bitter *et al*., 2024), reinforcing the notion that evolutionary and ecological processes can occur on the same timescale and potentially interact (Hairston *et al*., 2005; Carroll *et al*., 2007). One of the mechanisms by which populations change in response to environmental variation is adaptive tracking (Rudman *et al*., 2022; Bitter *et al*., 2024). In environments where resources are variable, and this variability creates distinct selection pressures, populations could potentially track shifts in resource abundance, and/or composition. What remains unknown is whether populations can adaptively track the distinct selective regimes associated with the fine-scale variation in environmental resources that shift over short timescales (e.g., weeks to months, 2-5 generations) and, if so, the extent to which such resource variation then simultaneously impacts both ecological and evolutionary dynamics in natural populations. Alternatively, variation in the resource environment may be associated with other evolutionary responses to environmental heterogeneity such as adaptive plasticity (Ghalambor *et al*., 2007; Charmantier *et al*., 2008; Brooker *et al*., 2022) or bet-hedging (Stearns, 1976; Gillespie, 1977; Simons, 2011).

Natural populations of *D. melanogaster* experience pronounced seasonal variation in the resource environment: in the ancestral range, the putative ancestral substrate marula (Mansourian *et al*., 2018) is seasonally ephemeral, as are the predominant fruit substrates in temperate regions (Reaume & Sokolowski, 2006; Markow, 2015). Seasonal environmental change, including variation in resource availability, affects population size (Dempster *et al*., 1995; Lawson *et al*., 2015), relative abundance of competing Drosophila species (Grainger *et al*., 2021), and patterns of rapid adaptation at both the phenotypic (Schmidt & Conde, 2006; Behrman *et al*., 2015; Behrman & Schmidt, 2022) and genomic (Bergland *et al*., 2014; Machado *et al*., 2021) levels. Similarly, experiments conducted in field mesocosms over the spring-to-fall seasonal progression have shown that *D. melanogaster* populations adaptively track shifts in the biotic and abiotic environment over weekly to monthly timescales (Rudman *et al*., 2019, 2022; Grainger *et al*., 2021; Bitter *et al*., 2024). However, the role of variation in the resource environment in contributing to the ecological and evolutionary dynamics of populations remains unresolved.

Dietary manipulations in the laboratory have revealed the direct influence of caloric and macronutrient variation on *D. melanogaster* life history traits, including viability, fecundity, body size, lifespan, and appendage size (Chiang & Hodson, 1950; Bakker, 1959, 1962; Wang & Clark, 1995; Skorupa *et al*., 2008; Rodrigues *et al*., 2015; Shingleton *et al*., 2017). Diet variation also affects patterns of genomic evolution (Kawecki *et al*., 2021) and multiple phenotypes, including mate choice (Schultzhaus *et al*., 2017) and ethanol tolerance (Cavener & Clegg, 1981). Natural populations are also characterized by standing genetic variation for macronutrient tolerance (Havula *et al*., 2022), suggesting populations could evolve rapidly in response to shifts in the resource landscape.

Here, we conducted a manipulative field experiment to examine the effects of diet on population dynamics and patterns of seasonal adaptation. Throughout a five-month experiment, we tracked census size, fitness-associated phenotypes, and allele frequencies genome-wide across replicate populations to determine how direct alterations of the nutritional environment impact ecological dynamics, phenotypic trajectories, and patterns of genomic evolution. We hypothesized that the dietary treatment would affect population demographic patterns and that nutritional variation might cause divergence in selective pressures, generating distinct patterns of adaptation across resource environments.

## METHODS

### Field Mesocosm Experimental Setup

Experimental populations were created using the same methodology as described in (Rudman *et al*., 2022). Briefly, a panel of eighty inbred lines originally collected in June 2012 (Linvilla Orchards, Media, PA, USA) were outcrossed and expanded for four generations of recombination. In the next generation, experimental flies were collected as a single 12h cohort from density-controlled cultures; collections were combined in groups of 2500 flies (equal sex ratios) and randomly assigned to an experimental treatment and replicate field cage (N = 6 cages per treatment).

Two distinct food substrates were developed to simulate unique resource environments. Our low-quality nutritional environment (LQ) is largely pureed apples and mimics a natural diet regularly experienced in temperate *D. melanogaster* populations. Alternatively, the high-quality nutritional environment (HQ) is a standard cornmeal molasses-based medium used in laboratory culture and prior experiments in this system (Rajpurohit *et al*., 2016; Rudman *et al*., 2019; Bitter *et al*., 2024). The principal differences between the nutritional treatments are caloric content and the ratio of protein to carbohydrates (recipes and nutritional information are given in Table S1).

Each population was maintained outdoors in a 2m^3^ mesh enclosure containing a fruitless, dwarf peach tree. The only food source and egg-laying substrate was 400ml of the respective dietary treatment contained in 900ml aluminum loaf pans, which were replaced every other day for the duration of the experiment (July 11th – November 23rd, 2020). After 2d of oviposition, each loaf pan was covered with a screen mesh lid and embryos were allowed to develop separately in a small cage until eclosion ceased. Loaf pans of media within experimental cages were protected from rain and direct sun on shelving units oriented away from direct sunlight.

### Measurement of population size and fitness-associated phenotypes

To observe how the nutritional treatments affected population dynamics, population size was estimated bi-weekly. At a standardized photoperiodic timepoint, 4 permanent ceiling quadrats in each cage were photographed. The number of adult *D. melanogaster* in each sample photograph was counted (using a modified cell counting protocol with ImageJ (O’Brien *et al*., 2016)) and corrected for total mesocosm surface area to estimate population size (Rudman *et al*., 2019, 2022).

In addition to population demographics, we phenotyped each population for a variety of fitness-associated traits, which have been shown to respond to variability in the seasonal and resource environments (Behrman *et al*., 2015; Behrman & Schmidt, 2022; Rudman *et al*., 2022). Our primary goal was to examine changes in the genetic composition of experimental populations over seasonal time and whether patterns of evolutionary change were altered by dietary treatment. Thus, all phenotypes at all timepoints were assayed in a common, laboratory environment. Eggs were collected overnight from each population on fresh loaf pans and returned to the lab for two generations of density-controlled, common-garden rearing (25°C, 12L:12D) on their respective diets. Fitness-associated phenotypes of the F2 generation were measured for each field cage on density and age-controlled replicates (N=3 per cage per timepoint). Fecundity was measured as the total number of eggs laid by a group of five females over three days (Paaby *et al*., 2014). The development time for males and females was estimated as time from oviposition to eclosion for groups of 50 eggs by recording eclosion events 3x per day at 09:00, 13:00, and 17:00 and scoring by sex (Behrman *et al*., 2015); viability was estimated as the percentage of eggs laid that emerged as adults. Starvation resistance was measured separately for males and females as the time to death for replicate groups of 10 individuals kept in vials on 1% agar. Thorax length was measured as the longest length across the dorsal shield in lateral view for 15 ethanol-preserved females. Measurements were recorded using a Leica MZ9.5 microscope, with an Olympus DP73 camera and CellSens standard measuring software (Betancourt *et al*., 2021). We assayed each of these phenotypes in the founding population five times during the experiment. All project data and analyses are available at https://github.com/jkbeltz/Nutritional_Quality_Analysis.git.

Temperature data were measured using a shielded, non-aspirated temperature/relative-humidity sensor (CS-215L) mounted at 2m height, located centrally among populations, and logged with a CR-1000 datalogger (all Campbell Scientific, Logan, UT). Temperature was logged every 15 minutes as an average of the previous 5 minutes. These data were used to construct degree day models (Kamiyama *et al*., 2020) to estimate generation times between each sampling interval throughout the experiment. 180-degree days were used to represent a single generation. Rates of phenotypic evolution were then calculated in haldanes (Gingerich, 2001) for all traits across the entire experiment as well as between each sampling interval.

In addition to the standard, common garden phenotyping of all cage populations at all timepoints on their diet of origin, an additional protocol was conducted at two timepoints (t2 on 18 September, representing the end of the summer period and expansion, and t4 on 25 October, before the winter collapse). Individuals from each population were reared on both dietary treatments to examine: 1) diet-associated plasticity for the five assayed phenotypes, and 2) whether such patterns of plasticity changed throughout the experiment. Eggs were collected overnight from each population on fresh loaf pans on their respective diets and returned to the lab, where individual eggs were manually removed and transferred to both diets. Fitness-associated phenotypes were measured on density and age-controlled replicates in the F2 generation following the same protocols described above.

Phenotypic data was analyzed in R (v4.3.2) using mixed-effect linear models, where time, treatment, and/ or phenotypic treatment were fixed values (Tables S2-S4), and individual cage/ sampling date was a random effect (lme4, fit by REML). T-tests, ANOVAs using the Satterthwaite method (Satterthwaite, 1946), and MANOVA were utilized to test the significance of phenotypic variation across treatments in multidimensional space (Table S6).

### Whole Genome Sequencing and Allele Frequency Estimation

We examined how the experimental manipulation affected patterns of allele frequency change over seasonal time. Specifically, during each of the five collection timepoints (Fig. 1), 100 F1 adult female flies were randomly selected from each cohort of field-collected eggs, preserved in 80% ethanol, and stored at –80° C. An additional four replicate samples were collected from the founding population used to initiate the experiment. Genomic DNA was later extracted from all pools using the New England Biolabs Monarch Genomic DNA Purification Kit. Whole-genome sequencing libraries were derived from each pooled sample with Illumina DNA Prep Tagmentation kit, followed by sequencing on the Illumina NovaSeq 6000 flow cell using 150-bp, paired-end sequencing reads. Raw sequencing reads were trimmed of adapter sequences and bases with quality score < 20 and subsequently aligned to the v5.39 *D. melanogaster* genome using bwa and default parameters (Li & Durbin, 2009). Deduplication of aligned reads was then conducted using Picard tools (http://broadinstitute.github.io/picard/). We downsampled aligned reads for each sample to obtain an equivalent, genome-wide coverage of 7x across samples. Haplotype-informed allele frequencies were then computed for each sample using the genome sequences of the founding inbred strains, a local inference method (Kessner *et al*., 2013) and a previously published pipeline (Tilk *et al*., 2019). This method produces an accuracy of allele frequency estimates at a genome-wide depth of 5x comparable to standard pooled sequencing approaches and 100x coverage (Tilk *et al*., 2019; Rudman *et al*., 2022). The final set of allele frequencies was then filtered to obtain only those single nucleotide polymorphisms (SNPs) with an average minor allele frequency > 0.02 in founder populations and >0.01 in at least one of the evolved samples, resulting in 1.9 M SNPs for our final assessment of genomic changes throughout the experiment.

**Figure 1.**
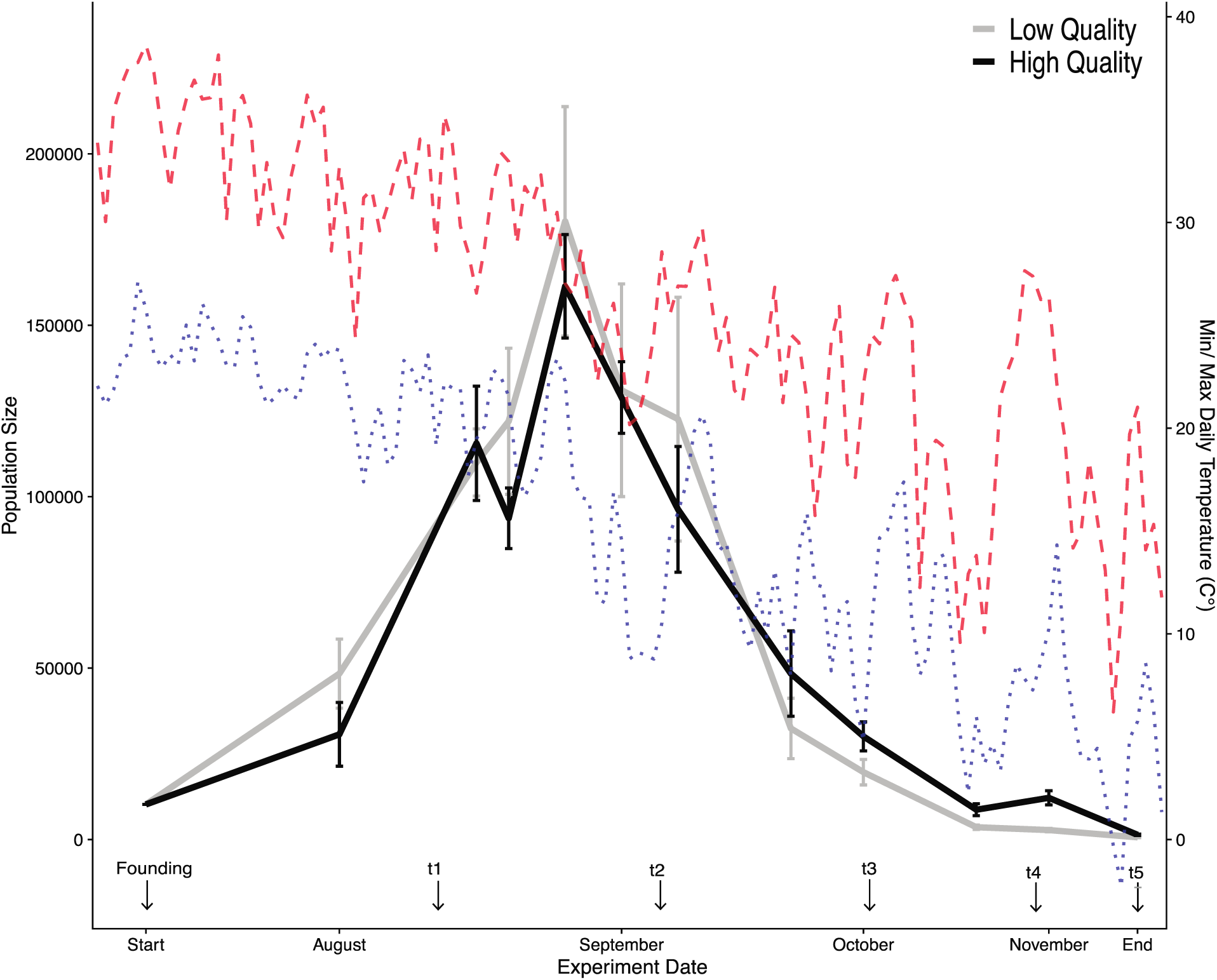
Mean (± s.e.) estimated census size across seasonally evolving populations of *D. melanogaster* reared in either a low-or high-quality nutritional environment, along with the daily minimum (blue) and maximum (red) temperatures throughout the experiment. Census estimates were obtained on 7/30, 8/12, 8/21, 8/30, 9/2, 9/18, 9/26, 10/1, 10/20, 11/2, and 11/14/20. The founding of the experimental populations and all phenotyping sampling timepoints are labeled on the x-axis.

### Analysis of Genome-Wide Allele frequencies

Analysis of genome-wide allele frequency data was conducted in R v. 3.5.6. We first quantified and visualized patterns of genomic variation across all collected samples using Principal Component Analysis (*prcomp* package). Allele frequencies were centered and scaled prior to PCA, and samples were projected onto the first, second, and third principal components. Through this, we interrogated whether variance in genome-wide allele frequencies throughout the experiment was predominantly driven by collection timepoint, treatment, or a combination thereof.

Next, we identified individual SNPs with systematic frequency movement across replicates throughout the experiment using a generalized linear model (GLM), first running this model independently for each treatment. As drift is unlikely to produce coordinated allele frequency across independent populations, we inferred that SNPs identified via this approach are subject to direct or linked selection in response to the shared seasonal selection pressures in our mesocosm. Prior to GLM regression, allele frequencies were weighted by effective coverage (Tilk et al. 2019), and the total number of chromosomes sequenced (N = 200). The normalized frequencies were fit to a GLM of the form: allele frequency ∼ timepoint, using a quasi-binomial error variance structure (Wiberg *et al*., 2017). P-values were adjusted using a Benjamini-Hochberg false discovery rate correction and the distribution of SNP significance as a function of chromosomal position was then visualized using Manhattan plots.

Next, we quantified the extent to which patterns of selection were parallel across treatments. Specifically, we reciprocally quantified the behavior of the rising allele at SNPs identified via GLM (FDR < 0.05 and allele frequency shift > 2 %) in each treatment independently in each replicate of the respective treatment from time points 1 to 5. We then compared each set of SNPs to a matched control set (matched on chromosomal arm and starting frequency) using a paired t-test to ask whether the magnitude of allele frequency movement exceeded background shifts at control sites and whether the dominant direction of movement was conserved in the test cage/treatment (FDR < 0.05). For visualization purposes, we plotted the median shift of target and matched control SNPs for each iteration of this analysis, separately for each chromosomal arm and genome-wide.

To further quantify treatment-specific patterns of allele frequency within specific chromosomal arms, we next used the GLM signal procured independently for each treatment and identified unlinked, genomic windows (hereafter, ‘clusters’) enriched in SNPs moving systematically throughout the experiment (method developed in (Rudman et al. 2022) and code provided: https://github.com/greensii/dros-adaptive-tracking).We then quantified the behavior of these unlinked clusters in the opposing treatment (e.g., the behavior of loci identified in HQ then examined LQ, and vice versa). Specifically, for each cluster, we computed the mean frequency shift of all seasonally evolving alleles (FDR < 0.1; effect size > 0.5%) in the opposing, ‘test’ treatment. To test whether the resulting distributions of allele frequency shifts indicated parallel change across treatments, we compared them to allele frequency distributions derived from sets of matched control SNPs (i.e., each target SNP within each cluster was matched to a control SNP located on the same chromosomal arm and within 5% starting frequency in the founding population).

Finally, we used another GLM-based approach to validate evidence of SNPs with different evolutionary dynamics between treatments and throughout the experiment produced via the approaches described above. Specifically, we used coverage and population-size normalized allele frequency data across all timepoints and treatments and ran a GLM of the form: allele frequency ∼ timepoint + treatment + treatment*timepoint. We then identified those SNPs with a significant treatment by time interaction as those with an FDR-corrected p-value < 0.1. We plotted the trajectories of these treatment by time SNPs for each treatment throughout the experiment and visualized their genomic distribution using Manhattan plots.

## RESULTS

### The ecological impact of variation in resource quality

Census estimates reveal that variation in the nutritional environment did not produce significant differences in population size across the two treatment groups, with both exhibiting similar patterns of seasonal growth throughout the summer, achievement of maximum size in early September, and subsequent autumnal decline (Fig. 1). There was no overall effect of treatment on population size across the experiment (Table S2a). Furthermore, no individual pairwise comparison between treatment groups was significant at any individual timepoint after multiple testing corrections (Table S2b). These results demonstrate that, contrary to our prediction, the resource quality variation imposed here does not impact adult census size in this system.

### The impact of nutritional variation on trait evolution

Our longitudinal phenotypic analysis revealed patterns of rapid evolutionary change across all surveyed traits (Fig. 2A-G, Table S3). The two dietary treatments had a pronounced and direct effect on phenotype, as expected: this is evident in the phenotypes exhibited by the founding population cultured on the two diets, as well as in the separation between treatments in the multivariate analysis (Fig. 2G). Our primary interest, however, was to examine whether patterns of evolution over seasonal time were parallel or divergent between the two dietary treatments, as demonstrated by a significant interaction between time and treatment. We find that two traits show distinct patterns of evolutionary change between treatments over seasonal time: viability and body size (Table S3). Viability remained roughly consistent over time for the LQ treatment; for the HQ treatment, viability remained constant until the last sampling interval, over which it declined by approximately 30%. Similarly, body size was generally constant over time for both treatments from t1 to t4. Over the last sampling interval, however, body size evolved to be larger in the LQ treatment while continuing to decrease in the HQ treatment. These differential patterns of phenotypic evolution over seasonal time were concentrated at the last timepoint, where rates of evolution were also observed to be noticeably higher as temperatures and population sizes declined.

**Figure 2.**
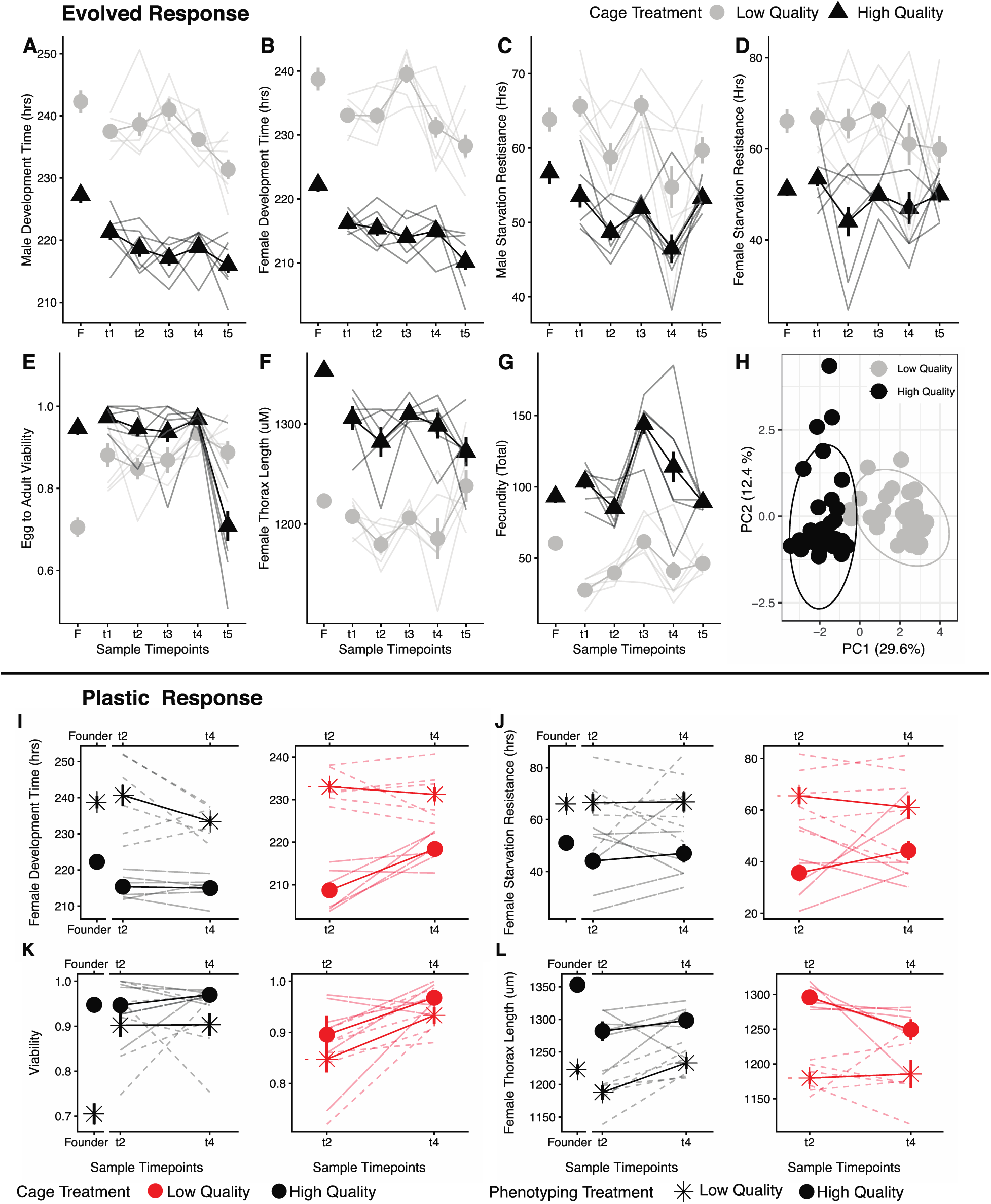
Patterns of phenotypic evolution (A-G) and phenotypic plasticity (I-L) over seasonal time in two distinct resource environments. Mean trait values (± s.e.) are depicted by bolded lines with symbols (LQ in grey, HQ in black), and individual replicate cages are given as non-bolded lines (A-G). Founder populations are included for presentation purposes only. Variation in the nutritional environment impacts all assayed traits and shifts patterns of seasonal adaptation for two traits (E,F). When all trait means across all time points are used to construct a principal component analysis (H), a strong effect of nutritional environment is observed with a significant effect in a MANOVA (F_2,58_=1.28E+02, p=2.2E-16). Patterns of phenotypic plasticity for HQ and LQ populations assayed on both diets at two timepoints (I-L; plastic response for all traits given in Fig S1). Mean trait values (± s.e.) are depicted by bolded lines with symbols (treatment denoted by color (LQ in red, HQ in black), and assay treatment denoted by shape (LQ denoted by an asterisk, HQ by solid points), and individual replicate cages are given as non-bolded lines. Founder populations are included for presentation purposes only. While trait value is affected by the assay environment, there is no effect of dietary treatment on the evolution of the plastic response (Table S5.)

At two timepoints (t2 & t4), all population trait values were assayed on both diets in a common garden laboratory environment. All traits exhibited pronounced plasticity in response to the dietary environment (Fig. 2I-L). However, we found no evidence that the treatment manipulation affected how nutritional plasticity changed over time (Table S5), as no three-way interaction (time x experimental treatment x assay diet) was significant for any of the measured phenotypes. For multiple traits, however, we did observe shifts in nutritional plasticity over time (viability, female development rate), suggesting that the evolution of plasticity may be an essential component of rapid adaptation to environmental change in this system.

Rates of phenotypic evolution in haldanes (H) were determined for all traits between each sampling time point for both treatment groups (Fig. S3a). Rates of phenotypic evolution increase systemically across time intervals, with the fastest rates observed in the final time interval (t4 to t5) for all phenotypes. Interestingly, when viewed across the entire experiment (t1 to t5), rates are lower; this is consistent with the observation that rates of evolution, even over the timescales examined here, are determined by measurement interval (Gingerich, 1983). We found no effect of the nutritional environment on the pace of evolutionary change (Fig. S3b, Table S4); thus, the dietary manipulation affected evolutionary trajectories but not rates of change over generational time.

When conducting an experimental manipulation of dietary sources, we would be remiss not to address the potential effects of the microbial community (Figs. S3-S). Regular sampling and 16s sequencing of the mesocosm food supply, as well as wild-caught and F1 common garden-reared adults (Methods found in Supporting Materials), revealed that LQ samples tended to hold more lactic acid bacteria (LAB) in lab culturing and greater acetic acid bacteria (AAB) in the field, relative to HQ populations that were more consistent across environments (Fig. S8).

Overall, the total bacterial abundance of the food substrate (Fig. S7) and diversity (Fig. S7) / community composition (Fig. S8, Table S7) of the *D. melanogaster* host populations were not significantly affected by treatment. While this level of sequencing resolution cannot exclude the impact of subtle variation of individual taxa, these results indicate that the observed population dynamics and patterns of evolutionary change reflect a host response to the dietary treatments and not direct or indirect effects associated with distinct microbial communities.

### Impact of nutritional variation on genomic evolution

PCA indicated that the variance in allele frequencies across all samples was primarily influenced by seasonal time and secondarily by the resource environment (i.e., the sample segregation along PCs 2 and 3 corresponded with the collection timepoint and resource environment, respectively, as shown in Fig. 3A-B). The appreciable force of seasonal varying selection was further exemplified by our regression-based analysis. Specifically, for both the HQ and LQ treatments, we quantified genome-wide signatures of selection across replicates whereby thousands of SNPs, across all chromosomal arms, exhibited parallel movement (FDR < 0.05) of relatively large effect (frequency change > 2%) throughout the experiment (Fig. 3 B&C).

**Figure 3.**
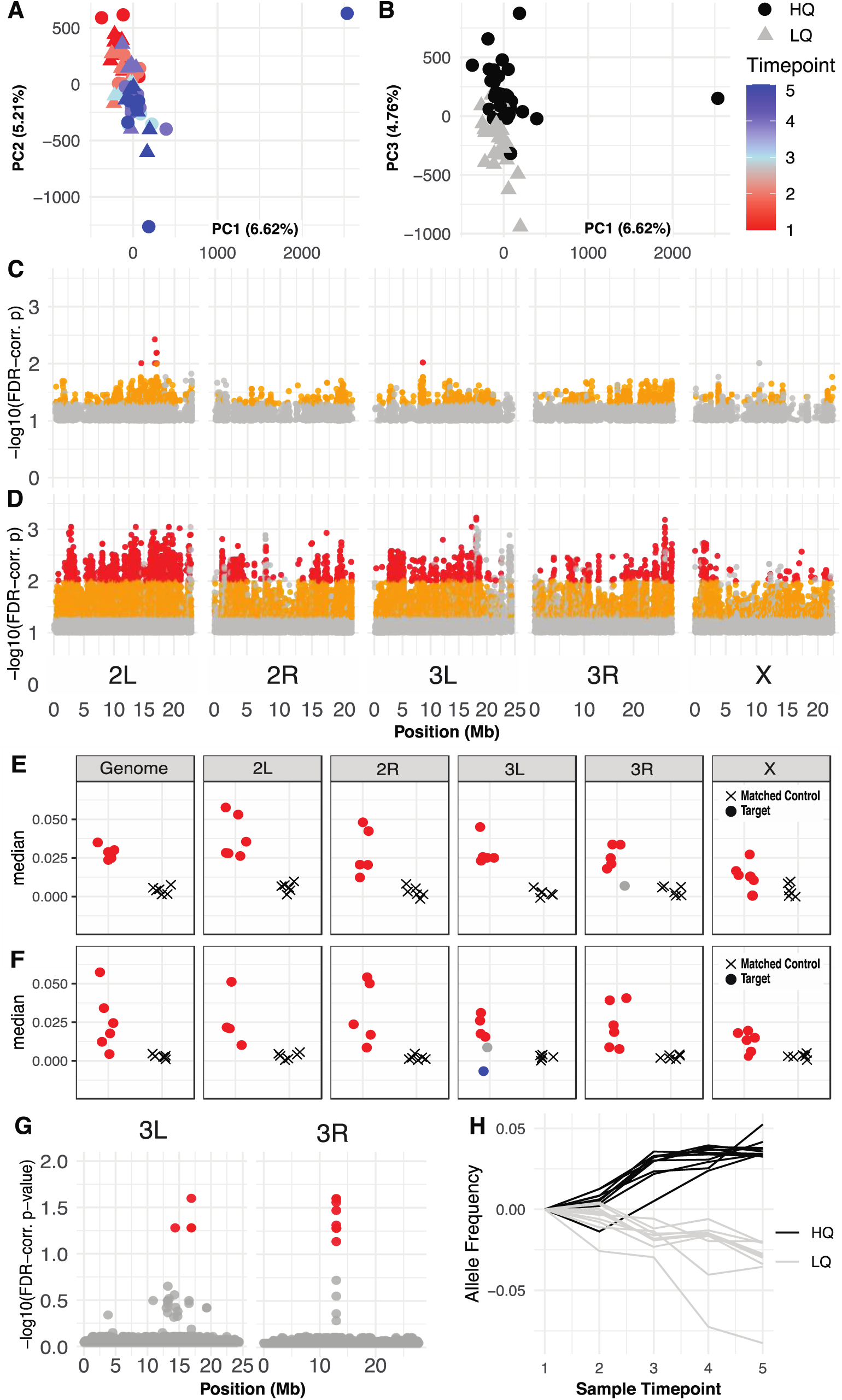
Genome-wide patterns of variation within and between treatments throughout the experiment. (A/B) Principal component analysis across all samples using allele frequency data from 1.9 million SNPs. In (A), samples are projected onto the first two principal components and colored by collection time point. MANOVA revealed a significant effect of time overall (F=32.65, p=5.07×10^-10^), as well as in the LQ (F=17.31, p=2.11×10^-5^) and HQ (F=3.61, p=3.16×10^-2^) populations individually, but no effect of treatment or the time by treatment interaction. Variation in PC1 is essentially driven by a single outlier sample, which is likely due to a more severe bottleneck in one cage over the t4 to t5 interval. In (B), samples are projected onto the first and third principal components and colored by treatment. Here, the MANOVA reveals a significant effect of treatment (F=25.88, p=1.31E-08, as well as time (F=5.21, p=8.49×10^-3^). C/D: Manhattan plots depicting SNPs shifting in parallel across replicates through time in LQ (C) and HQ (D). Point color corresponds to FDR-corrected p-value and effect size (grey: FDR < 0.1 and effect size > 0.5%; orange: FDR < 0.05 and effect size > 2%; red: FDR < 0.01 and effect size > 2%). E/F: Cross-treatment validation for seasonally evolving SNPs (identified in B/C). In each treatment, seasonally evolving sites (FDR < 0.05, effect size > 2%) were identified using all six populations and were then compared to the shifts at those sites in the alternative treatment populations. The set of shifts at the rising allele was compared to shifts at background sites matched for chromosome and initial frequency. Plotted is the median shift for the matched controls (Circles), and test sites (Xs), which are colored red if the test sites were significantly parallel (red), blue if anti-parallel (blue), and grey if there is no significant difference from the background movement (gray). SNPs were identified in the LQ treatment and tested in the HQ treatment populations (E), genome-wide and within each chromosome arm, and were also identified in the HQ and tested in the LQ populations (F). (G) Manhattan plot of alleles that shift differentially over time between treatments (FDR < 0.1; only chromosomal arms 3L and 3R depicted, as no significant SNPs were identified on the remaining chromosomes). Fourteen SNPs were identified with a significant treatment by time interaction via a GLM (red points). (H) Mean frequency trajectories of significant SNPs depicted in (G) throughout the experiment, separately for the HQ (black) and LQ (gray) replicate cages.

Given the observed differences in phenotypic evolution (Fig. 2) and the indication that genome-wide allele frequencies were perturbed between treatments (Fig. 3A-B), we next aimed to specifically assess the degree of parallelism in the genomic response to seasonal changes between HQ and LQ replicates. To achieve this, we measured the behavior of seasonally evolving SNPs (FDR < 0.05 and allele frequency change > 2%) identified independently within each treatment (Fig. 3B&C) in all replicate cages of the opposing treatment throughout the course of the experiment (Fig. 3 E&F). This analysis revealed that the response to seasonal changes at the muller element level is predominantly parallel, whereby the primary direction of allele frequency movement is concordant between treatments with a magnitude of change that exceeds of background allele frequency movement (red points, Fig. 3E&F). Interestingly, this parallelism was reduced on 3L and 3R and, in one instance, was antiparallel (blue point, Fig. 3F), providing evidence that a subset of loci may exhibit patterns of divergent seasonal responses between treatments.

We next sought to quantify the relative contribution of parallel vs. anti-parallel seasonal selection between treatments across the genome. To do this, we used the set of seasonal SNPs identified independently for each treatment (Fig. 3 B&C) to compute the minimum number of putatively unlinked loci (hereafter, ‘clusters’) associated with these signals of selection (Methods). This analysis revealed 81 clusters in HQ and 117 clusters in LQ. We then quantified the behavior of the rising allele at all GLM-significant SNPs within each cluster in the opposing treatment relative to a set of matched controls. Our findings again showed predominantly parallel shifts between treatments, with 96% of clusters in HQ and 95% in LQ exhibiting significantly parallel behavior in the opposing treatment. Notably, however, we identified one cluster in HQ and three clusters in LQ that demonstrated antiparallel movement between treatments (Fig. S4), indicating that selection was acting in opposite directions. As suggested by the analysis in Fig. 3 (E&F), this antiparallel movement was confined to chromosome arms 3L and 3R. Overall, these results illustrate a largely parallel genomic response to seasonal changes across treatments, with distinct deviations from parallelism consistently observed.

Finally, we aimed to identify individual SNPs with treatment-specific trajectories. To achieve this, we conducted an additional GLM of the form: allele frequency ∼ time point + treatment*time point. This model identified many SNPs with systematic movement across all samples through seasonal time (e.g. 16,776 SNPs at an FDR < 0.1) and 14 SNPs that displayed a significant treatment-by-time interaction (FDR <0.1). Ten of the treatment*time SNPs are located on chromosome 3R (positions 12959498 – 12998860), while four are on 3L (one at position 14360958 and three others between positions 16911479 – 16911529; Fig. 3G, Table S6). This result underscores the extent of the shared seasonal response compared to the relatively localized divergent selection between treatments over time. Visualization of the trajectories of these 14 SNPs throughout the experiment illustrates their treatment-specific dynamics.

Specifically, each SNP demonstrated a magnitude of allele frequency change greater than ∼3%, with the direction of allele frequency movement being opposite between treatments in all cases. These patterns indicate a set of loci under strong selection in both nutrient environments but selected in opposing directions (Fig. 3H). Collectively, the genomic data revealed pervasive seasonal evolution within each treatment, as well as evidence of localized divergent responses to selection between treatments. Therefore, we find that distinct resource environments contribute to differing patterns of evolution at both the genomic and phenotypic levels.

## DISCUSSION

We report the observations from a large-scale, replicated field experiment in which the dietary resources of half of the populations were significantly reduced (20% fewer calories, 60% reduction in protein). Despite this difference in dietary resources, the largely parallel response within and between treatments indicates the robustness of the population response to seasonally dynamic environments. However, we also show that the trajectory of phenotypic evolution is altered by dietary treatment, as are patterns of allele frequency change at multiple regions of the genome. Given the parallelism among replicates in patterns of evolutionary change, we interpret these patterns as adaptive: diet impacts how experimental populations adapt to the changing seasonal environment.

Our dietary treatments elicited very distinct phenotypic responses in all populations, which manifests prominently when surveyed on the alternative diet. The HQ diet is a typical rich diet for laboratory culture and the diet used in prior mesocosm field experiments in this system (Bergland *et al*., 2014; Rudman *et al*., 2019, 2022; Grainger *et al*., 2021; Bitter *et al*., 2024). The LQ diet was designed to mimic a natural, late-season diet in temperate environments. Natural populations of *D. melanogaster* in temperate North America persist on a wide range of substrates that vary in nutritional/caloric content and are seasonally ephemeral as well as spatially variable (e.g., strawberries, citrus, cherries, grapes, persimmons, paw paws, peaches, pears, apples, etc.). The results we present here suggest that variation in the dietary environment can result in distinct selection pressures that may contribute to spatially and temporally variable selection and, in turn, to the maintenance of genetic variation in natural populations.

Ecologically, we hypothesized that resource quality would impact population dynamics, whereby the LQ replicates would exhibit a limited rate of growth and reduced carrying capacity relative to the HQ treatment. This was rejected, as no distinction between treatments was observed in population growth patterns or maximum population size. Several potential explanations exist for these results. Our experimental populations may, in fact, not have been resource-limited by our experimentally imposed diets. However, all available food (400 ml) was consumed in 14 days, and mesocosm populations’ reproductive output during summer is ∼10,000–30,000 eggs per cage every 2 days (Rudman *et al*., 2022). Additionally, dietary treatment could have influenced rates of senescence and average lifespan, making population sizes similar across treatments despite different underlying age structures. Although diet affects lifespan in *D. melanogaster* (Skorupa *et al*., 2008; Zanco *et al*., 2021), estimates (Lee *et al*., 2008) suggest similar lifespans for both treatment groups. Treatment differences in the intensity of intraspecific competition might also explain this result: HQ populations, with higher fecundity and viability, could initially surpass LQ populations until growth is limited by competition (Connell, 1983), yet LQ census sizes exceeded HQ throughout the first two months. However, as reproductive output in the field was not surveyed directly, potential effects of differences in larval density across treatment cannot be excluded. Furthermore, differences in substrate crowding due to treatment effects on body size (flies were ∼10% larger on HQ) may limit HQ population growth due to increased competition. Lastly, our dietary manipulation might have altered the microbial community, leading to indirect host effects. We found no significant differences in bacterial abundance or notable taxonomic differences between treatments, indicating that microbial-derived protein or mutualistic effects are likely equivalent across treatments. Variations in the microbial community appear insufficient to impact long-term population demographics. Regardless, our results demonstrate that a ∼20% increase in the energy input does not influence the rate of population growth, maximum size, or persistence. We speculate that a combination of factors may contribute to this, but overall, the ecological dynamics in our system are likely primarily driven by intra-specific competition that may be modulated in multiple ways by the nutritional environment.

Evolutionarily, variation in the nutritional environment did not restrict the capacity, rate, or magnitude of the adaptive response of populations; while the response to seasonal shifts across all populations was generally robust, the dietary manipulation did cause divergence in evolutionary trajectories for a subset of phenotypes (viability and body size) and had a quantifiable impact on patterns of allele frequency shifts over the course of the experiment. The treatment divergence, both genomically and phenotypically, demonstrates that these populations can adaptively track and respond to shifts in the resource environment, as well as the seasonal environment, concurrently. We identified 14 SNPs that demonstrated opposite patterns of allele frequency change between treatments. Over half of the SNPs with treatment-by-time interactions are in a ∼40kb window on chromosome 3R, within a region associated with inversions (3RP/3RK) that have been linked to phenotypic and adaptive divergence (Durmaz *et al*., 2018; Kapun *et al*., 2023). The remaining SNPs were found on 3L in two regions separated by 2.5 Mbp. These 14 SNPs do not represent 14 independent loci of selection: given the recombinational landscape in *D. melanogaster* (Comeron *et al*., 2012; Hunter *et al*., 2016) and the composition of the founding population, we showthat this is likely driven by selection on three genomic regions with strong linkage disequilibrium in each. Overall, the divergence in the selection landscape between treatment groups suggests that the genetic architecture underlying this treatment effect is relatively simple, which stands in stark contrast to the highly polygenic seasonal adaptive response.

The highest rates of phenotypic change and the greatest divergence in trait evolution between treatment groups (Fig. 2D, E, F & 3C) were observed between the final sampling timepoints when temperatures dipped below freezing. This suggests that selection pressures may be most pronounced during this pre-winter phase of the season, magnifying the effect of our dietary treatments. Similarly, increased rates of evolutionary change during this period (post-frost) have been previously observed (Bergland *et al*., 2014; Grainger *et al*., 2021; Behrman & Schmidt, 2022; Bitter *et al*., 2024), suggesting that this relatively punctuated selective landscape may be critical in establishing seasonal cycling of both phenotypic means and allele frequencies (Schmidt and Conde 2006; Bergland et al. 2014; Behrman and Schmidt 2022). The dynamics of overwintering and seasonal adaptation in *D. melanogaster* populations may also be strongly impacted by intense selection in the pre-winter phase, and variation in the resource environment may be especially consequential during this period (Saunders *et al*., 1990; Kimura *et al*., 1992; Kimura & Yoshida, 1995; Huang *et al*., 2021).

We observe a significant impact of dietary resources on the evolutionary paths of populations, with no observable effect on ecological outcomes. This finding challenges the traditional notion that divergence among populations in fundamental ecological parameters (e.g., census size) are evident on shorter timescales than evolutionary change, and indicates that evolution can transpire without altering at least some metrics of population ecology.

Consequently, this suggests that interpretation of ecological metrics should consider underlying evolutionary divergence that may not manifest in demographic outcomes. Moreover, this implies that monitoring evolutionary changes may serve as an effective measure for detecting short-term population shifts, as our findings hint that it can reveal subtle alterations with greater sensitivity.

*D. melanogaster* populations can adaptively track seasonal heterogeneity (Rudman *et al*., 2022; Bitter *et al*., 2024), and we now report a simultaneous adaptive response to an additional axis of environmental variation. The adaptive and divergent responses to variation in food supply may also affect patterns of mate choice (Schultzhaus *et al*., 2017) and other behaviors associated with food supply (Rosenberg *et al*., 2018). This ability to track and respond to fine-scale variation may drive the maintenance of genetic and phenotypic diversity globally, as natural populations respond to multiple axes of variation concurrently.

## SUPPLEMENTAL MATERIALS

### Microbial Sampling and Sequencing Methods

Since the Drosophila-associated microbiota shifts over space and time (Chandler *et al*., 2011; Wong *et al*., 2013), affects host phenotypes (Chandler *et al*., 2011; Wong *et al*., 2014; Keebaugh *et al*., 2018), and varies with dietary substrate (Massey *et al*., 2024; Yakovleva *et al*., 2024), we examined the host and environmental microbiota in all populations at two timepoints. On September 18 and October 25, approximately 100 adults were directly sampled by aspiration from each cage for microbial sequencing. Additionally, a sample of older food (which has been in the environment for at least 10 days, and still had flies actively eclosing) was collected from each cage to assess the microbial community associated with each dietary substrate. These samples, along with common-garden F1s from phenotyping samples, were stored at –80°C; upon completion of the experiment, DNA was extracted from three replicate pools of five male and five female flies, and amplicon libraries were generated using 16s rRNA V1V2 primers (Forward: 5’-AGAGTTTGATCCTGGCTCAG-3’, Reverse: 5’-TGCTGCCTCCCGTAGGAGT-3’) and Q5 polymerase before being paired-end sequenced with Illumina MiSeq to 15x coverage at the University of Pennsylvania / Children’s Hospital of Philadelphia microbiome core, following previously described protocols (López-Aladid *et al*., 2023). All reads identified as Wolbachia sp. were excluded from the analysis. All sequence analysis was performed using QIIME2 (following code provided at https://github.com/jbisanz/16Spipelines) and visualizations generated with R (version 4.3.2). Raw sequence data are available at https://github.com/jkbeltz/Nutritional_Quality_Analysis.git.

**Table S1:**
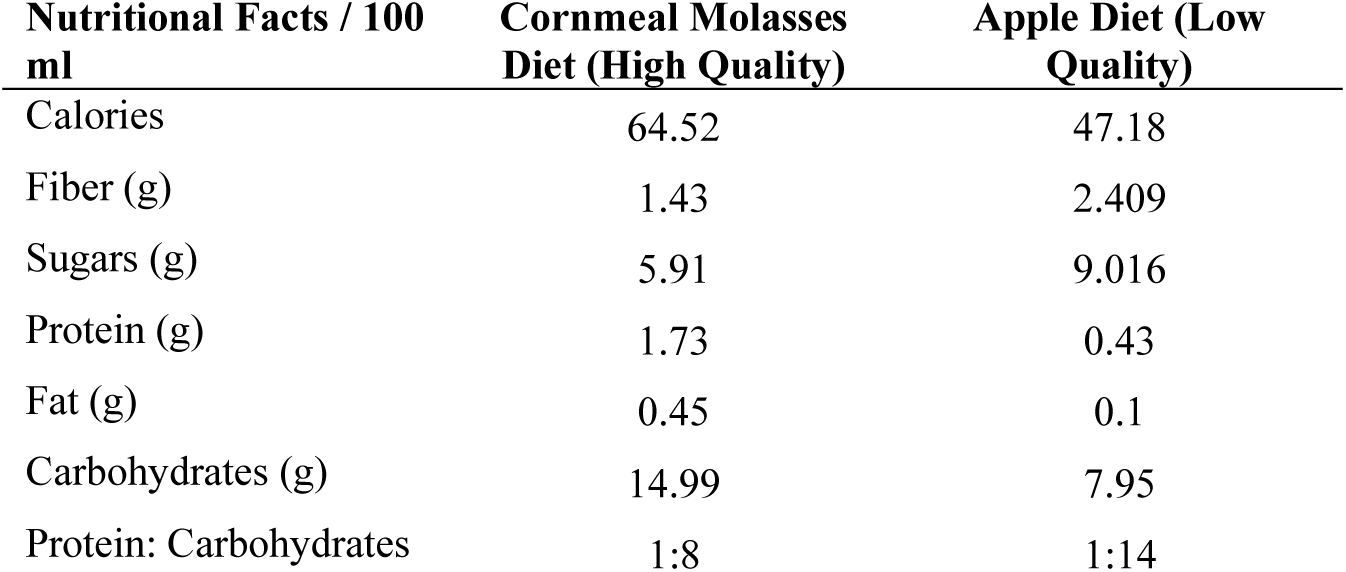
Nutritional composition for both dietary treatments. The lab standard diet (HQ) consists of 81.0% distilled water, 8.0% molasses (Golden Barrel Sulfur-Free Black Strap Molasses), 5.3% cornmeal (Flystuff 62-100), 3.0% ethanol, 1.5% inactive dry yeast (Flystuff 62-108), 0.73% soy flour (Flystuff 62-115), 0.5% agar (Flystuff 66-103), and 0.17% methylparaben (Apex Bioresearch Products 20-258) by weight. The semi-natural apple-based diet (LQ) consists of 55.3% unsweetened applesauce (Regal #10 Can Unsweetened Applesauce), 39.0% distilled water, 3.0% ethanol, 1.5% inactive dry yeast (Flystuff 62-108), 0.65% agar (Flystuff 66-103), and 0.17% methylparaben (Apex Bioresearch Products 20-258) by weight. Preparation of both the LQ and HQ diets followed the same cooking protocol. Macronutrient calculations were done using the Drosophila Dietary Composition Calculator (https://brodericklab.com/DDCC.php).

**Table S2 a & b.**
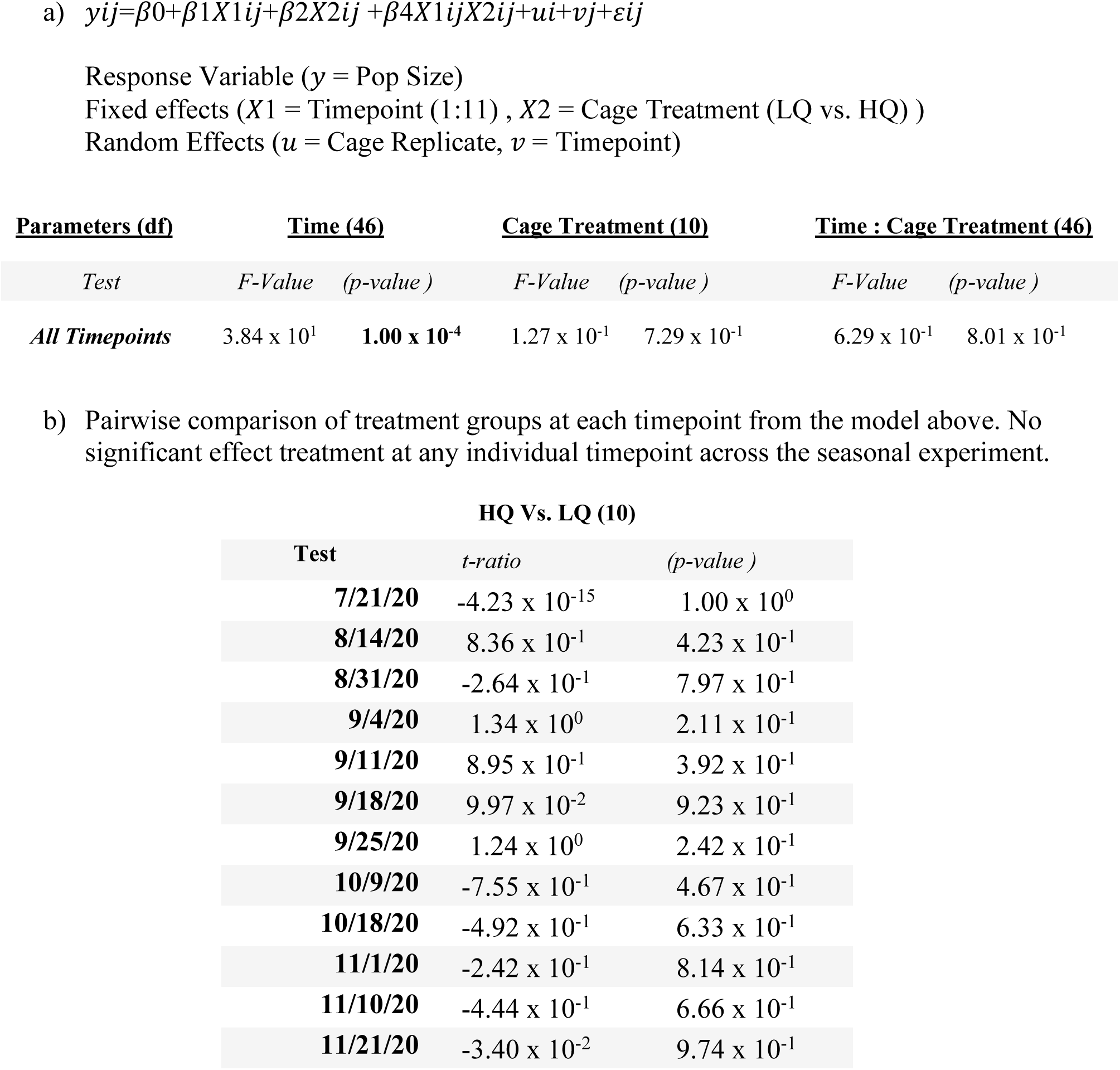
A linear mixed-effects model was constructed to determine the effect of nutritional quality on census size across replicate mesocosms. The model was tested across all 11 timepoints, and pairwise comparisons were made across the treatment group at each timepoint. ANOVA was used to assess the impact of sampling time and treatment on census size across all phases of the experiment. The model was constructed in R using lme4, and ANOVA was used for post hoc analysis. Linear mixed model fit by REML. t-tests use Satterthwaite's method [‘lmerModLmerTest’]

**Table S3:**
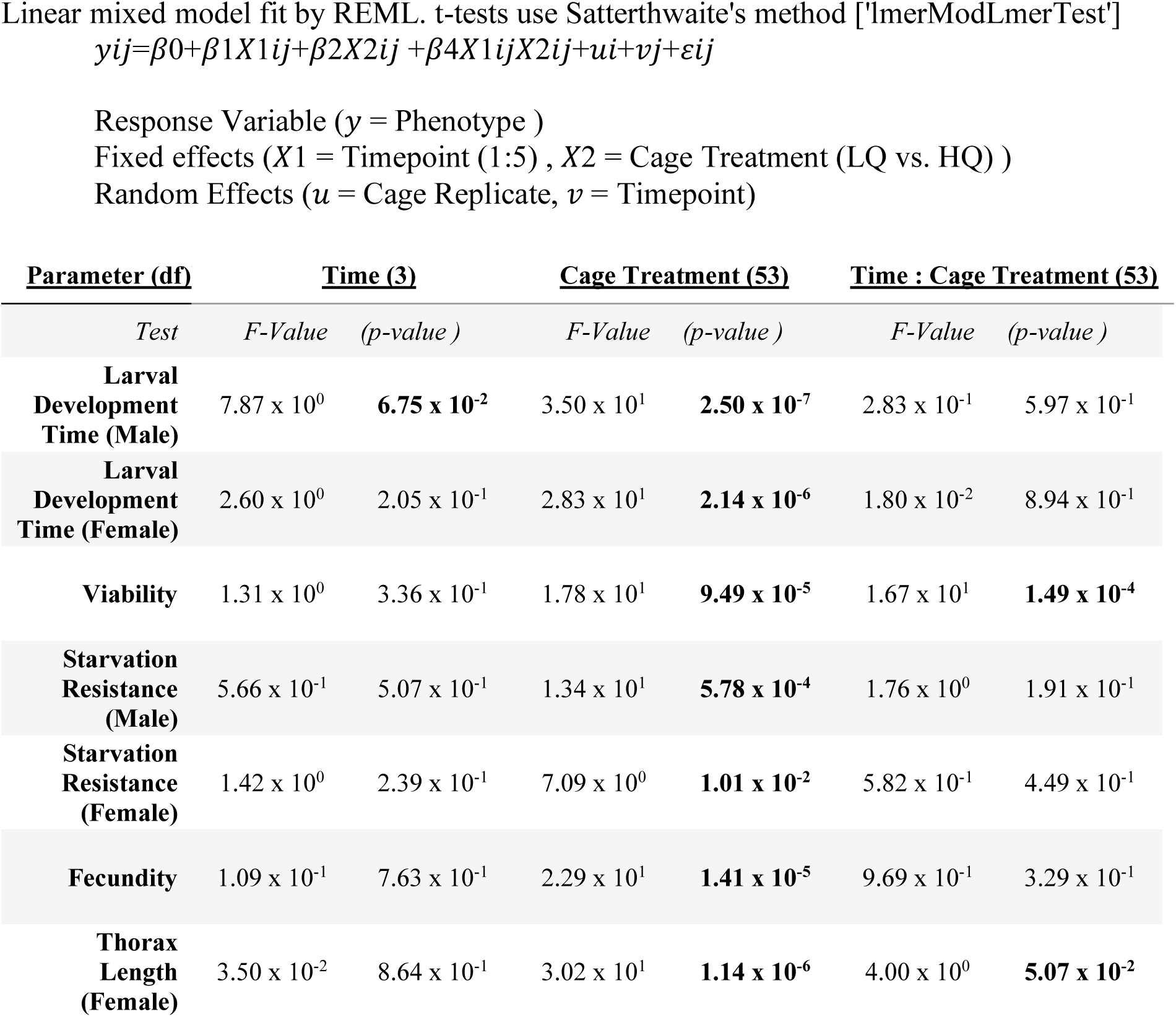
A linear mixed-effects model was constructed to determine if treatment (LQ & HQ), season (t1-5), or the interaction of both has effects on life history traits. Trait value is the response; treatment, timepoint, and the interaction term are predictors (individual cage treated as a random effect). Models were constructed for each trait individually in R using lme4, and anova() was used for post hoc analysis.

**Table S4 a & b.**
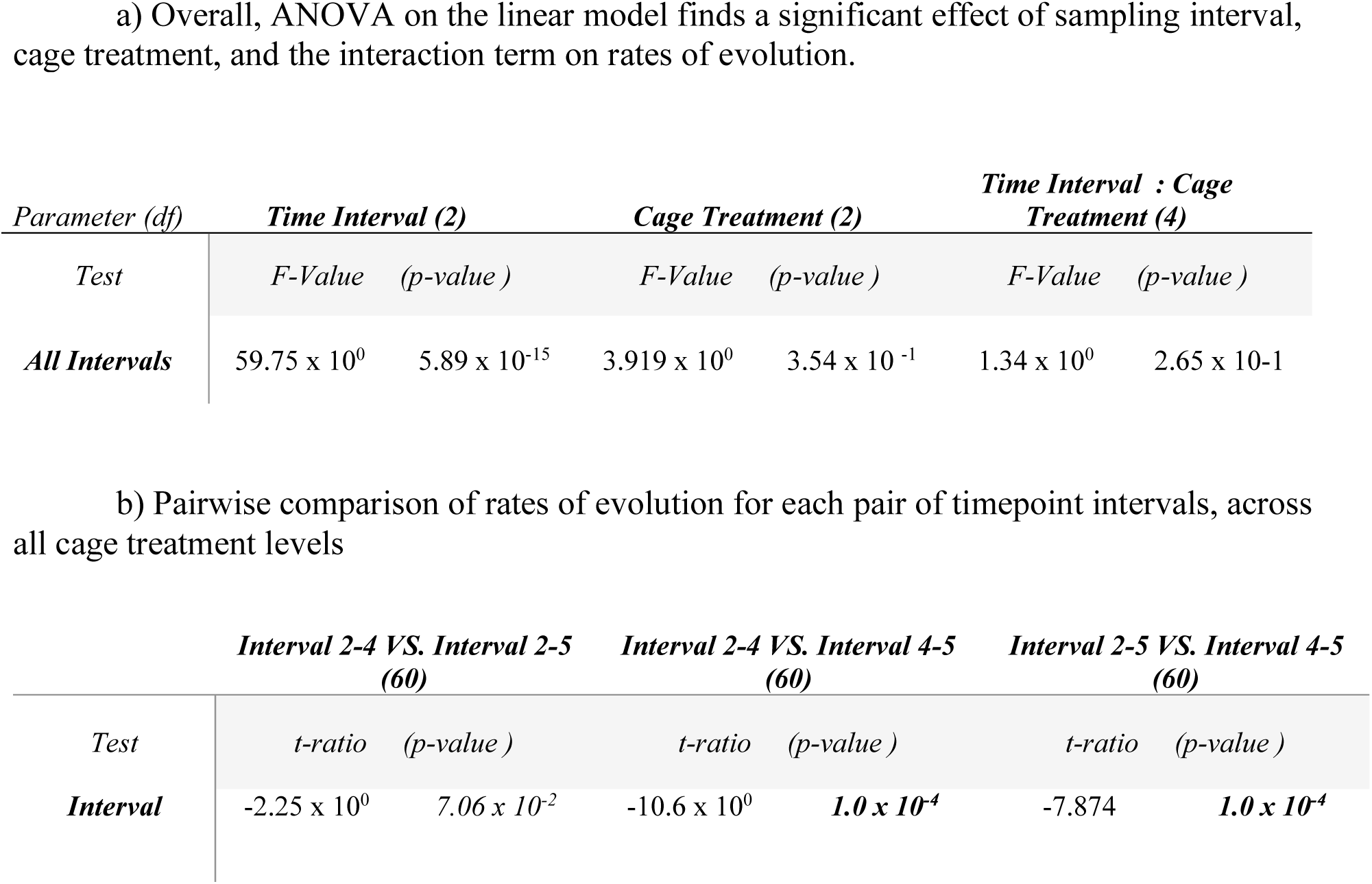
Linear model constructed to determine the effect of resource quality treatments on rates of evolution in haldane’s, for 8 life history phenotypes across 3 timepoint sampling intervals (2-4, 2-5, 4-5).

**Figure S1.**
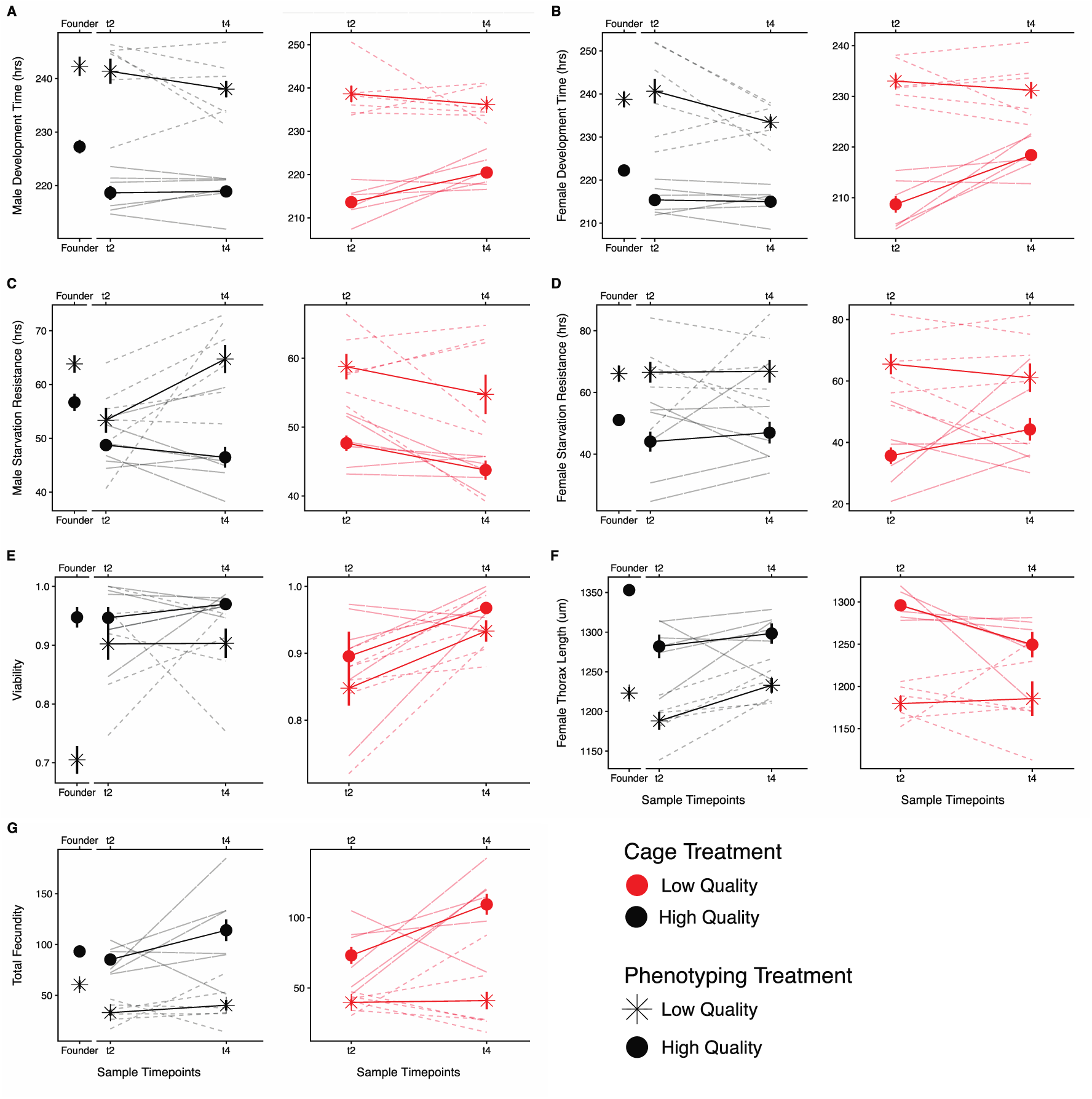
Mean (± one s.e.) phenotypes for replicate populations (N=6) reared on the low-quality (LQ, red) and high-quality (HQ, black) nutritional environments and assayed on LQ (asterisk/dash) and HQ (solid) environments at two experimental time points (mid-September(t2) & early November (t4)). Thinner dashed and solid lines represent individual replicate populations. Assaying each population in both treatment environments allowed us to determine if the plastic response to diet is evolving differentially across treatments and/or over seasonal time. The founder population, which was reared on the HQ diet, was also assayed in both environments, and is included purely for presentation purposes. We find that across treatments and time, assay diet primarily determines trait value. The mixed-effect linear model (Table S5) found that the phenotyping treatment (assay diet) significantly impacted all mean trait values, and some traits also showed an interaction between assay diet and time. We find no indication that the plastic response is evolving differentially across treatment groups and over time.

**Table S5:**
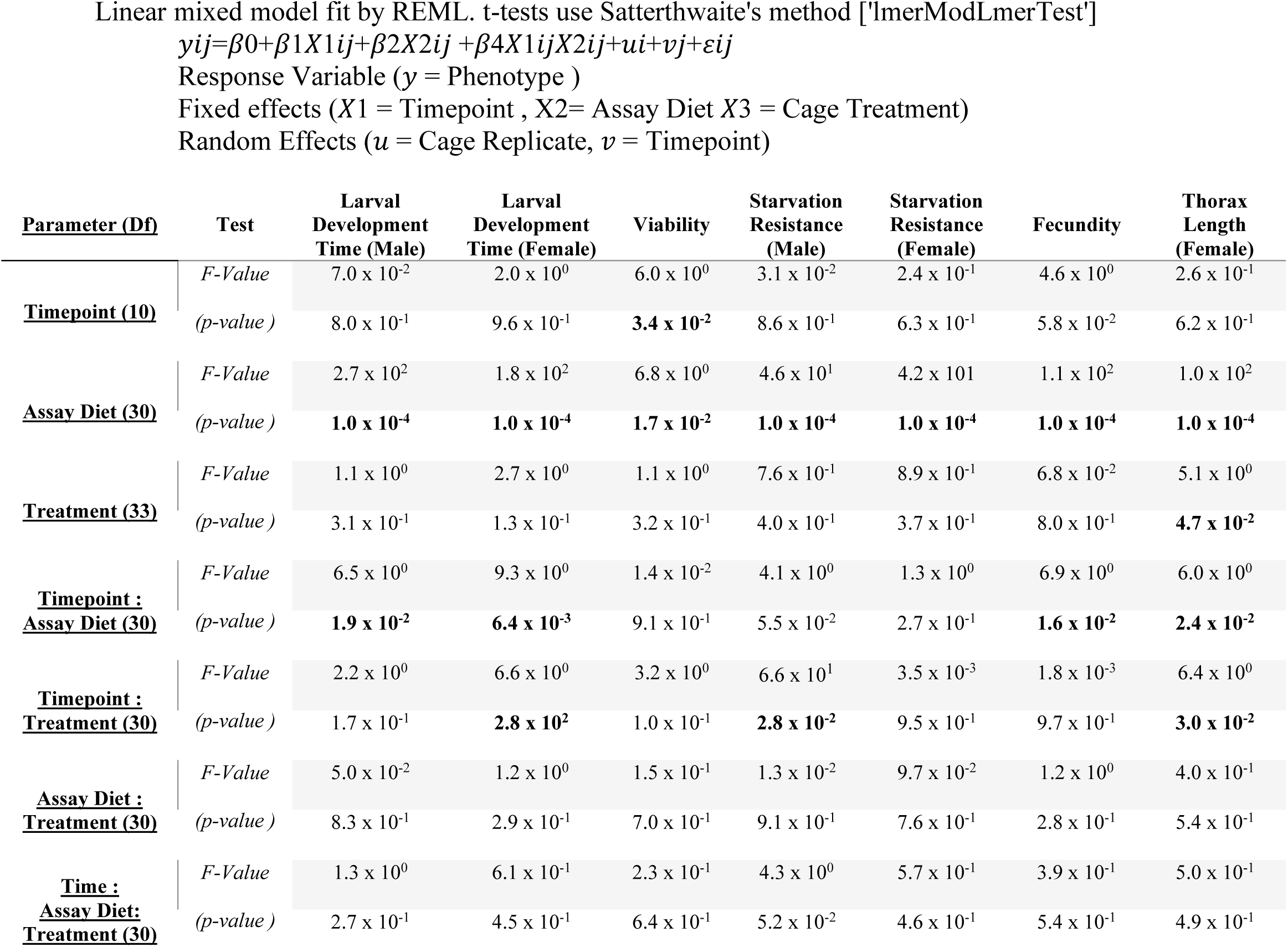
A three-way linear mixed-effects model was constructed to determine whether treatment (LQ vs. HQ), seasonal time (t2 vs. t4), assay diet (LQ vs. HQ), as well as any interaction terms, explain the patterns of variation across all life history traits. The model treated individual cage values and sampling timepoint as random effects. This model was constructed for each trait variable, and an ANOVA post hoc was conducted on each trained model. F-values and P-values are reported, and significant P-values are bolded. The model was constructed in R using lme4, with post hoc testing using anova(). As described in Fig S1, trait values were strongly driven by assay diet and the interaction of time and assay diet. Very little consistent effect of treatment was detected, and there is very little explanation offered by the interaction terms, showing that the plastic response is not evolving differentially across treatment and timepoint.

**Figure S2.**
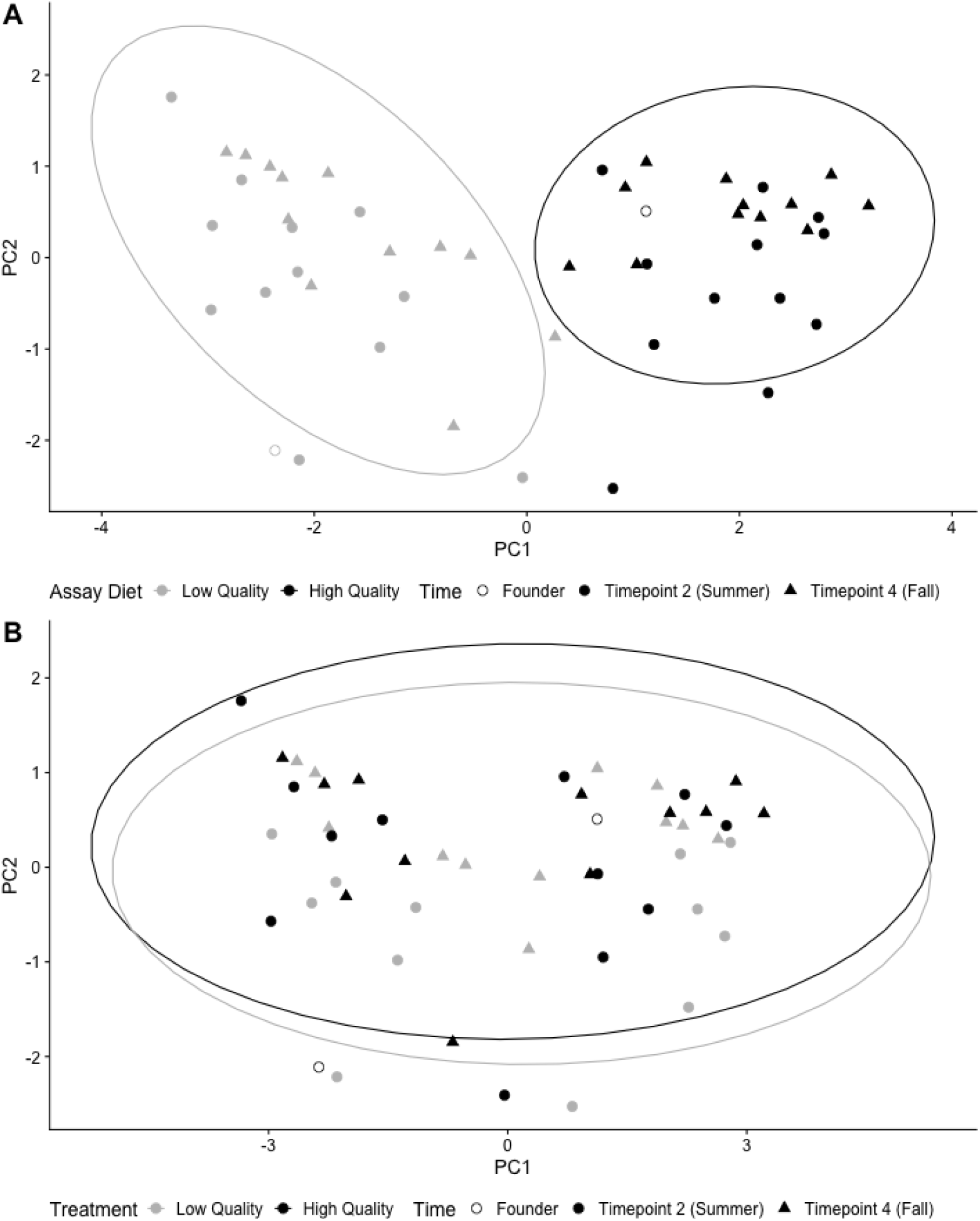
Principal component analysis was constructed using mean trait values across seven assayed traits (Fig. S1) of replicate populations (N=6) of *D. melanogaster*, reared in two distinct resource environments, collected at two timepoints, and assayed in both dietary environments. When points are shaded by the assay diet (A), we see a clear separation and a significant effect of the assay diet (F2,47=1.23E+02, p=2.2E-16). When the same points are shaded not by assay diet but by mesocosm treatment (B), we see complete overlap in the groupings (F2,47=6.36E-01, p=5.34E-01). This further demonstrates that when populations are assayed in both respective environments, the assay environment drives the trait values and that the plastic response to resource variation is not evolving differently across treatments.

**Figure S3.**
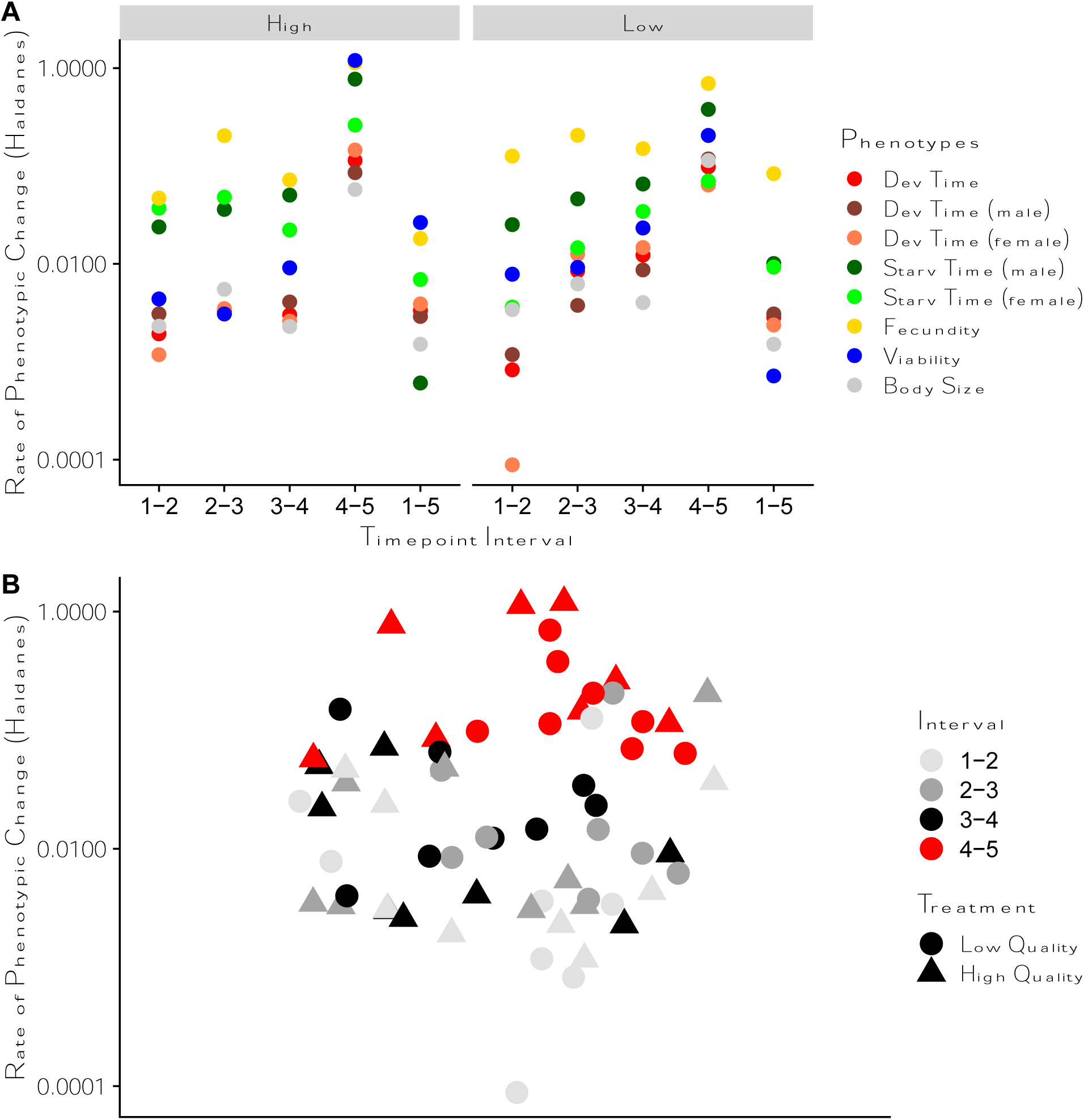
Rates of phenotypic evolution in Haldanes were calculated for all traits between each sampling interval. (A). When traits are combined (B), there is no significant impact of nutritional treatment (Table S4) on the rate of phenotypic change. The highest rates of trait change are consistently observed in the final time interval, where rates approach a standard deviation change per generation. The number of generations per time interval was estimated using a degree day model (Methods) and was determined to be 3.15 (Founder – t1), 2.28 (t1-t2), 1.26 (t2-t3), 1.3 (t3-t4), and 0.10 (t4-t5).

**Fig S4:**
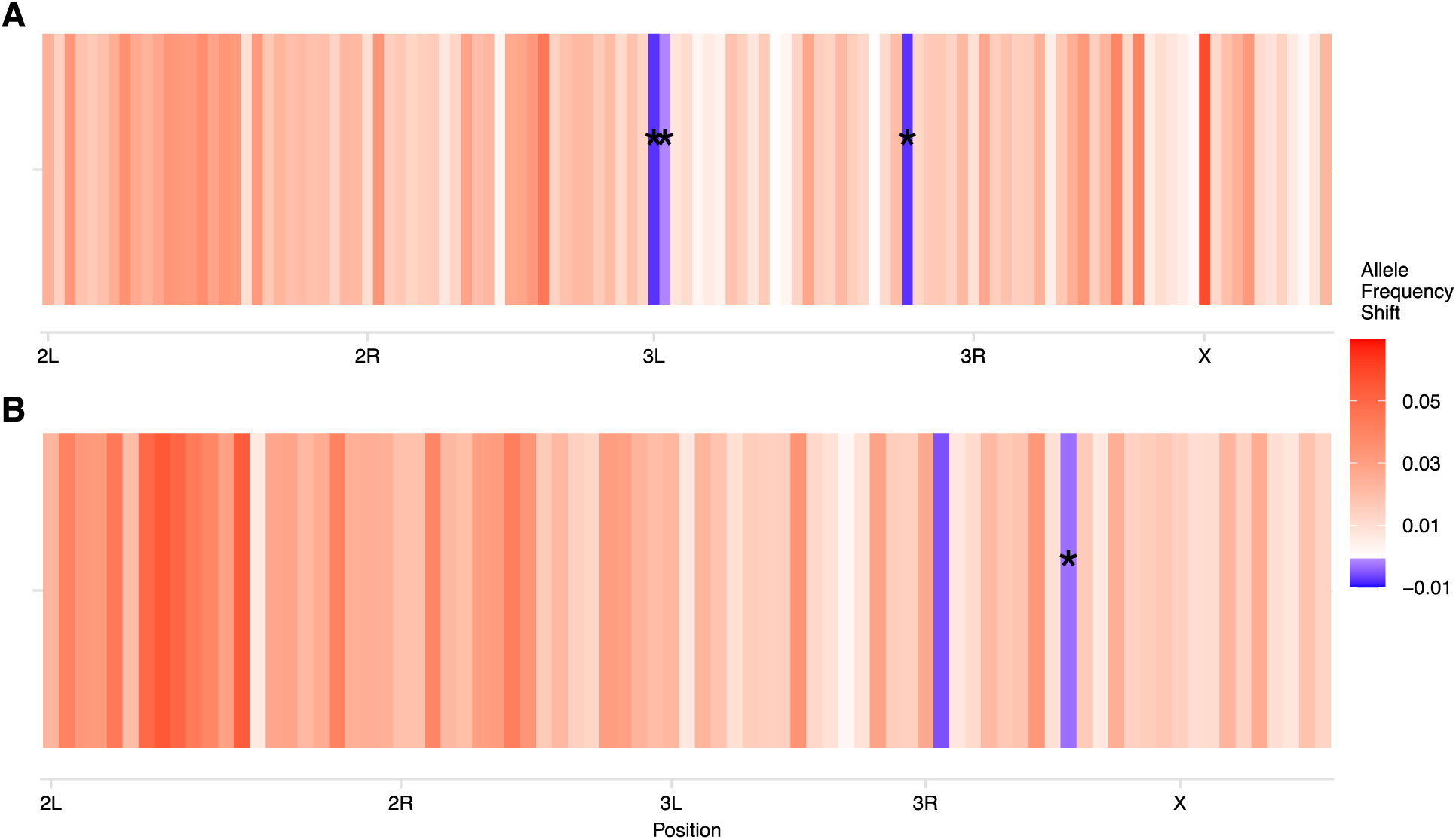
Quantification of cross-treatment parallelism using clusters of unlinked loci enriched in SNPs identified in Fig. 3. (A) corresponds to clusters identified in HQ and tested within LQ, while (B) corresponds to clusters identified in LQ and tested in HQ. Each bar corresponds to an unliked cluster, which is shaded by the magnitude of allele frequency change quantified in the opposing treatment. Colors describe the level of parallelism between clusters on a spectrum from red to white to blue, with the darkest red (shift > 0.03) indicating significant parallelism and blue (with an asterisk) a significant antiparallel relationship between the allele frequency response. White shades indicate neutrality (no change compared to background movement), and median shades of red and blue describe the relative parallelism / antiparallelism at these clusters, which have borderline or insignificant relationships.

**Table S6:**
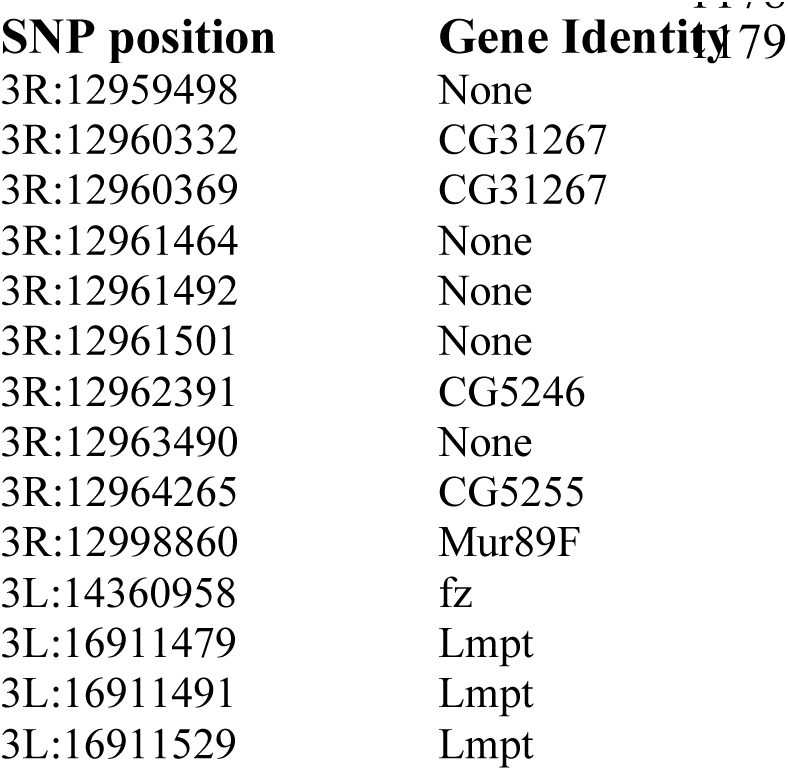
SNPs and genes associated with divergent selection between the HQ and LQ treatment groups. These SNPs showed a significant interaction between time and treatment in the GLM. Locations are listed in *D. melanogaster* genome version r5 dm3.

**Figure S5.**
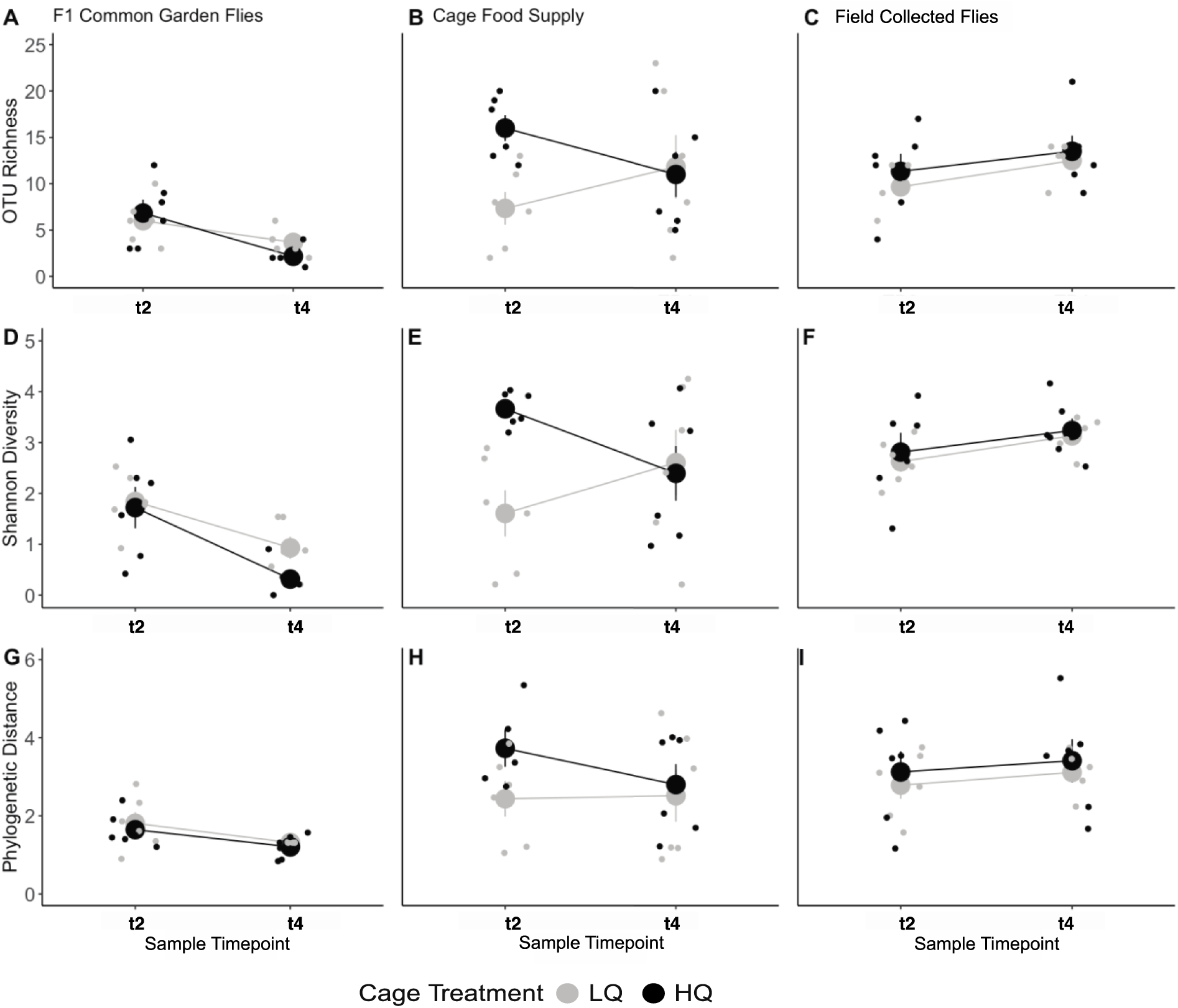
Alpha diversity metrics for bacterial communities in all populations and timepoints. Bacterial richness and diversity (Shannon & Faith_PD) for field-collected adults, food supplies, and lab-reared F1s at t2 (summer-evolved) and t4 (fall-evolved). Seasonality drives all alpha diversity metrics in field samples and F1-reared adult flies, with no significant separation in treatment means at either timepoint. For the cage food supply, we see greater richness and diversity in the HQ treatment in t2 and a convergence in t4, relative to diversity/richness in the LQ treatment. The nutritional manipulation had no effect on the microbial diversity of the host in either the field or the lab. This suggests that observed patterns of phenotypic and genomic evolutionary change are likely driven directly by nutritional manipulation and not an indirect effect of microbial diversity. Furthermore, the fact that the manipulation altered the food supply with no effect on bacterial associations suggests that the host largely controls the composition of its microbiome (top-down) as opposed to being dictated by the environment (bottom-up).

**Figure S6.**
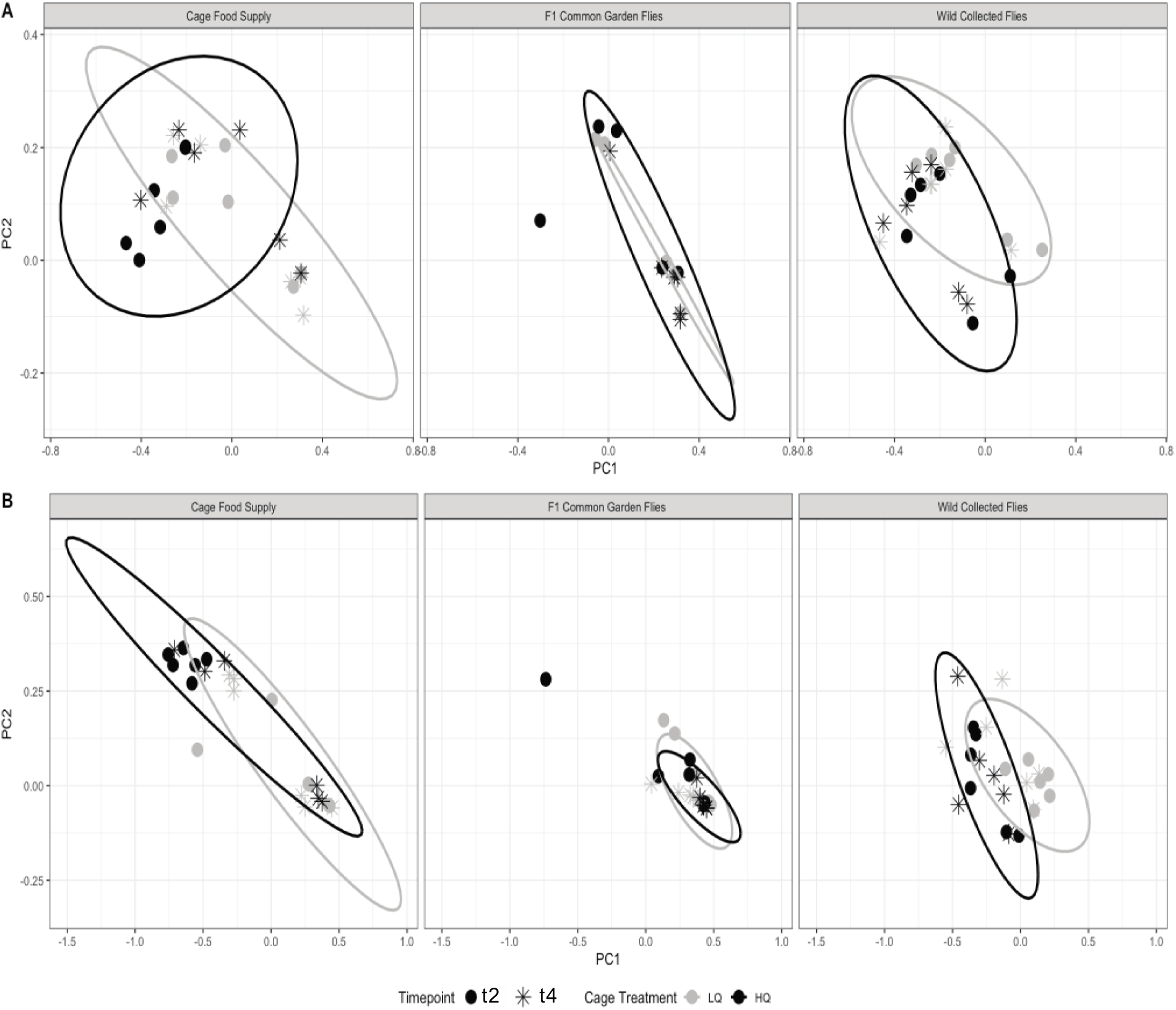
Principal component analysis was constructed using the relative abundance of all nonsingleton bacterial taxa in unweighted (a) and weighted (b) unifrac distance. Sequence variants were identified in field collected flies, cage food, and F1 flies reared in a single generation in a lab common garden, at experimental t2 (Summer Evolved, solid dots) and t4 (Fall Evolved, asterisks) for both LQ (grey) and HQ (black) populations. The MANOVA for significance in centroid position found no effect of nutritional quality on bacterial community composition and no effect of treatment, or the interaction term, for either unifrac distances in all samples.

**Table S7:**
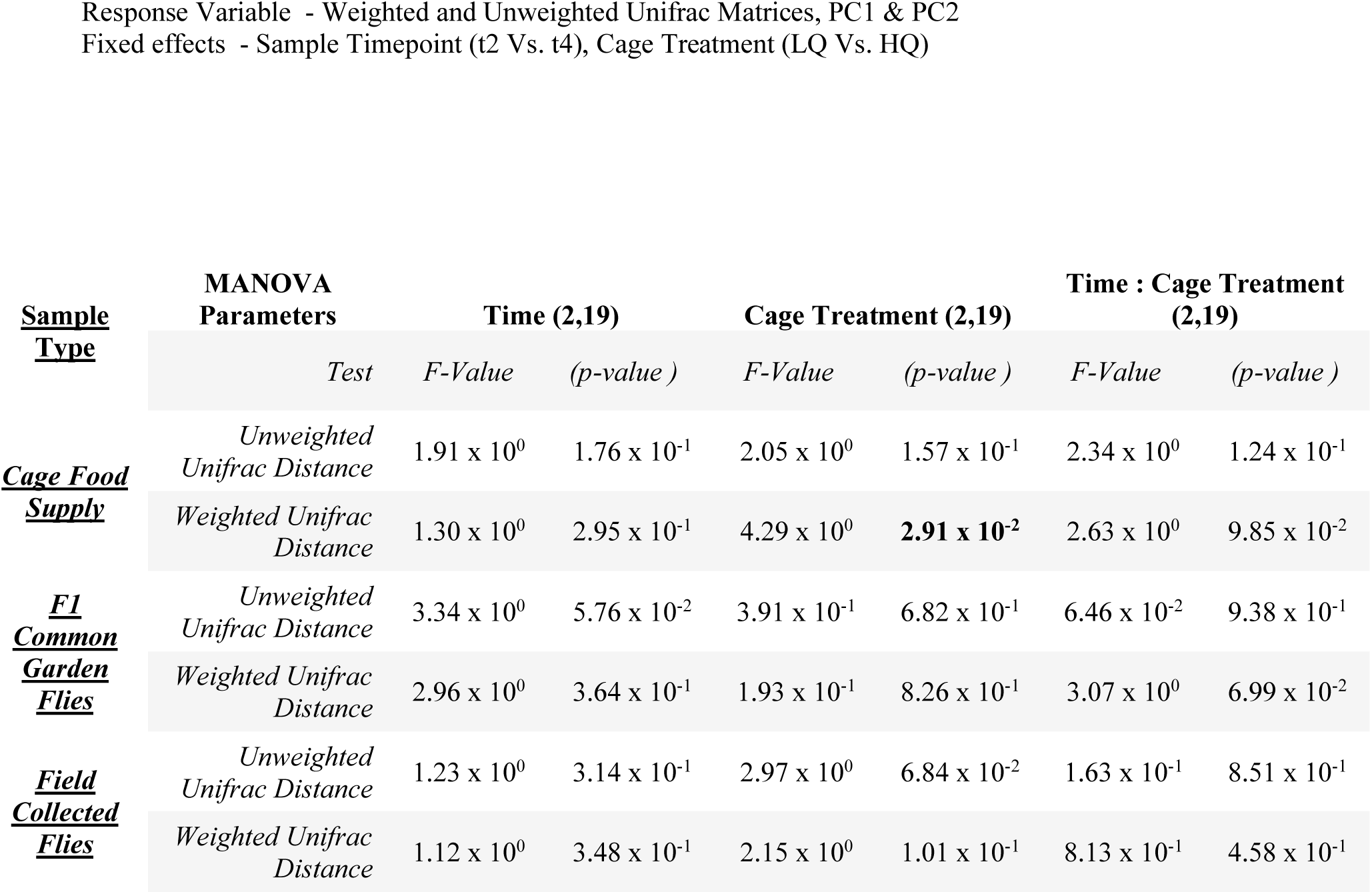
To assess whether the variance in the bacterial communities is significant between treatment groups for all sample types, we conducted MANOVA tests on the first two axes of variation for the weighted and unweighted Unifrac distance matrices constructed using the relative abundance of all non-singleton sequence variance. Like the alpha diversity metrics, the only clear effect on the microbial communities was observed in the cage food supply samples, which were significantly shaped by the treatment group. The bacterial communities of field-collected and F1 common garden flies showed no effect from cage treatment, time, or the interaction term. These findings again suggest that the ecological and evolutionary effects observed due to the treatment are not driven by variation in the microbial communities between treatments.

**Figure S7.**
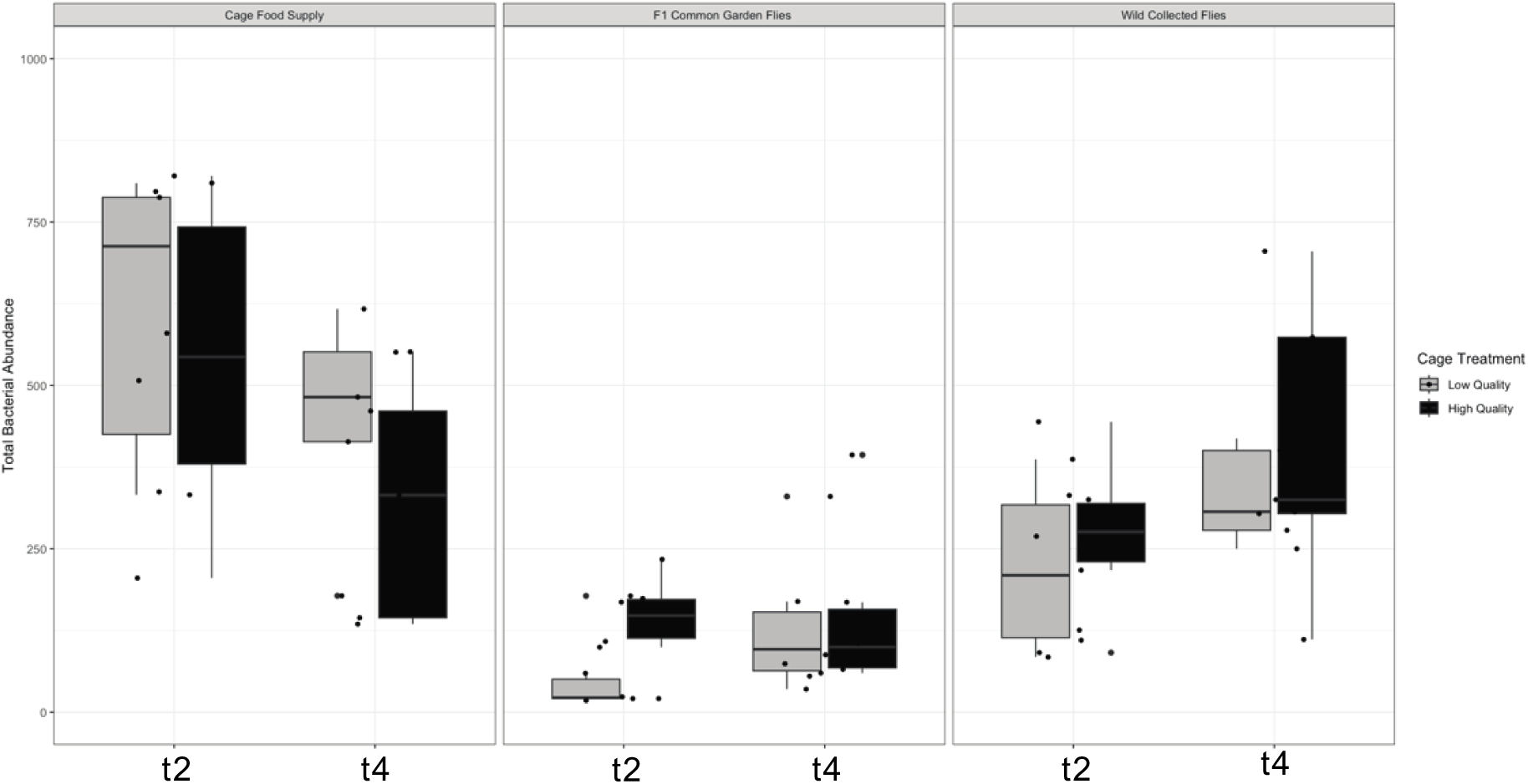
Total abundance of bacteria per sample across treatment, season, and sample type. Sequence variants were identified in wild-collected flies, cage food, and F1 flies reared for a single generation in a lab common garden, at experimental t2 (Summer Evolved, solid dots) and t4 (Fall Evolved, asterisks), for both Low Quality (LQ, Grey) and High Quality (HQ, Black) populations. The sequencing procedure and analysis protocol can be found in the methods. Raw counts of bacterial reads per sample were adjusted for the DNA concentration determined immediately after extraction to account for differences in DNA volume per sample. Seasonal change from summer to fall causes a decline in bacterial abundance in the food supply and an increase within wild-collected adults. T-tests revealed no significant effect of treatment on total bacterial abundance for wild adults and food at either time point. However, the HQ F1 common garden flies consistently have a greater abundance of bacteria than the LQ populations (p=4.21E-02). This result suggests that field treatment may impact how fly hosts take up bacterial taxa when brought back to a lab setting. Critically, the lack of effect of treatment on total bacterial abundance reveals that the volume of bacterial cells present on the food substrate is consistent across the two dietary treatments.

**Figure S8.**
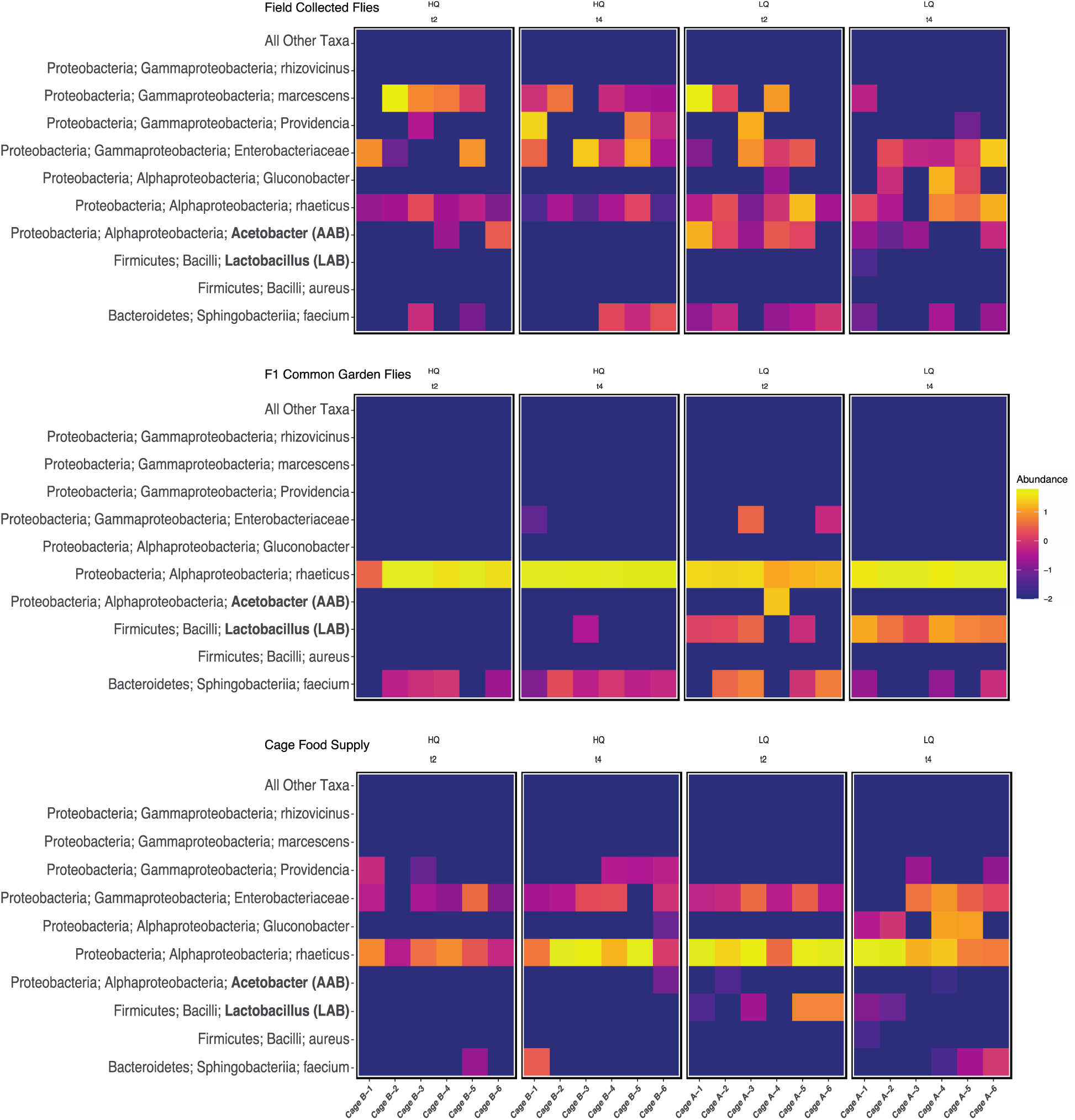
This heatmap illustrates the relative abundance of the ten most abundant bacterial taxa identified within and around the 12 experimental populations of D. melanogaster raised in one of two distinct nutritional environments. Major microbial groups are represented as rows, with each replicate cage as a column. HQ treatment samples are displayed on the left, while LQ samples are on the right, organized by timepoint (t2 and t4). Bacterial sequence variants were identified from samples of field-collected flies (top panels), cage food (middle), and flies raised for a single generation in a common garden laboratory environment (bottom). Bacterial sequence variants are classified by Phylum, Class, and then the lowest taxonomic groups identified with that variant (primarily Genus and species). The sequencing process and analysis protocol can be found in the methods. We observe a generally high level of consistency in the most dominant microbial taxa across treatment groups and time, with two notable exceptions. First, Lactobacillus are virtually absent in the field-collected flies from both treatments and timepoints, as well as in the HQ treatment samples from the food supply and F1 laboratory flies. In contrast, Lactobacillus are more abundant in the LQ treatment samples from the cage food and F1 laboratory flies at both timepoints. Second, field-collected flies from the LQ treatment show a higher prevalence of Acetobacter taxa than those collected from the HQ treatment, a pattern that is consistent across the two sampling timepoints. Interestingly, Acetobacter is found at low abundance and prevalence in the F1 and cage food samples at both timepoints. We also observe a greater prevalence of Gluconobacter taxa in the LQ populations, especially in the later sampling timepoint. As expected, we see clear differences between field and lab host samples (Ludington & Ja, 2020). When LQ and HQ samples are brought into a standardized lab environment, flies from 10 out of 12 LQ cages acquire Lactobacillus, while only 1 of 12 from HQ do the same. This suggests that the fly populations are evolving differently between dietary treatments and are “filtering” microbes in distinct ways. Overall, the total bacterial communities are not significantly different between treatments when flies are sampled directly from the field. However, given the level of sequencing resolution employed here, it is impossible to exclude the potential effects of treatment and seasonal evolution on individual bacterial taxa and strains.

